# Sustained interferon exposure creates a hyper-metastatic subset of melanoma cells

**DOI:** 10.64898/2026.04.01.715921

**Authors:** Mark H. Mannino, Tao Wei, Daniel Cassidy, Benjamin Knight, Devon L. Moose, Adrienne Shannon, Elena Piskounova, Zhiyu Zhao, Sean J. Morrison

**Affiliations:** Children’s Research Institute and Department of Pediatrics, University of Texas Southwestern Medical Center, Dallas, Texas 75390, USA; Department of Surgery, Surgical Oncology, University of Texas Southwestern Medical Center, Dallas, Texas 75390, USA; Department of Pathology, Beth Israel Deaconess Medical Center. Boston, MA, 02215; Howard Hughes Medical Institute, University of Texas Southwestern Medical Center, Dallas, Texas 75390, USA

## Abstract

We generated interferon signaling reporters in human and mouse melanoma cells and observed heterogeneity in interferon responses among cells in the same tumors. This was marked by inflamed regions within primary tumors that contained increased numbers of interferon-expressing macrophages/monocytes and elevated type I interferon signaling in melanoma cells. Melanoma cells that expressed GFP-Interferon stimulated gene 15 (ISG15) or GFP-Interferon-induced protein with tetraticopeptide repeats 3 (IFIT3) fusion reporters exhibited a profoundly increased ability to form metastatic tumors as compared to GFP-ISG15^-^ melanoma cells from the same tumors, particularly when transplanted into immunocompetent mice. The increased metastatic potential of GFP-ISG15^+^ cells was driven partly by increased CD47 expression, which protected metastasizing cells from phagocytosis by macrophages. Macrophages are thus a double-edged sword, inhibiting the development of metastatic disease by phagocytosing disseminated melanoma cells, but promoting the emergence of a hyper-metastatic subpopulation of cells in inflamed regions of primary tumors as a consequence of sustained interferon production.

**One sentence summary:** Sustained interferon exposure within inflamed regions of primary tumors dramatically increases the metastatic potential of a subpopulation of melanoma cells, partly by promoting CD47 expression, which protects against phagocytosis by macrophages.

---

Acute interferon exposure can inhibit cancer progression by promoting immune cell function^1–5^ and increasing the immunogenicity of cancer cells^6–9^. Conversely, chronic interferon exposure can promote cancer progression by exhausting T cells and promoting immune evasion by cancer cells^1,10–14^. Consistent with this, interferons are an important determinant of the response of cancer cells to immune checkpoint inhibitors^1,12^. Cancers that respond to immune checkpoint inhibitors tend to exhibit intermittent or acute interferon signaling^15^ while prolonged interferon signaling confers resistance by exhausting T cells^11^ and by increasing PD-L1 expression, which inhibits T cell function^10,14^. Lymph node-metastatic melanoma cells selected by serial passaging exhibit chronic, interferon-dependent epigenetic remodeling and increased PD-L1 expression, leading to improved lymph node colonization and subsequent expansion of regulatory T cells, which ultimately enable distant metastasis^16^. While these mechanisms enable metastasis to lymph nodes, they do not initially promote distant metastasis as lymph node-tropic melanoma cells are not more likely than control cells to form tumors after intravenous injection^16^. Finally, chromosomal instability and chronic cGAS/STING activation can promote metastasis through interferon-independent^17,18^ and interferon-dependent mechanisms^19^.

These differences in response to interferon signaling have typically been characterized among genetically distinct cancers (e.g. cancers with or without genomic instability) or cancers from different patients^20–22^. There is much less insight into differences in interferon signaling among cancer cells within the same tumor or the extent to which this confers differences in metastatic potential. Inflammation is not uniform within tumors, as immune hubs or other inflamed regions within tumors exhibit increased levels of inflammatory cytokines^23–25^, including interferons^26,27^. Primary melanomas contain hubs of PD-L1^hi^/HLA-A^hi^ melanoma cells in proximity to immune cells^26^ and patches of Interferon Regulatory Factor 1(IRF1)^high^ melanoma cells^28^ which are enriched at the invasive border, near macrophages and dendritic cells^27^.

Despite the heterogeneity in inflammation within tumors, few studies have compared the functions of cancer cells that differ in exposure to inflammation within the same tumor. An interferon-γ reporter in ovarian cancer cells showed that interferon-γ from T cells heterogeneously activates type II interferon signaling in certain regions of tumors^29^; however, it is unknown whether this is associated with functional differences in the ability of cancer cells to contribute to disease progression. We explored this question in melanoma by generating interferon signaling reporters that allowed us to compare the metastatic potential of melanoma cells from the same tumors that differed in their expression of interferon-regulated genes.

## Interferon-regulated gene expression in metastasizing cells

To identify mechanisms that regulate melanoma metastasis, we subcutaneously transplanted DsRed- and luciferase-tagged patient-derived melanomas (M405 and M481; see Extended Data Fig. 1a for known driver mutations in these melanomas) into NOD/SCID IL2Rγ^null^ (NSG) mice, then allowed the cells to form subcutaneous tumors and to spontaneously metastasize. When the subcutaneous tumors reached 2 cm in diameter, we killed the mice and flow cytometrically sorted DsRed^+^ human melanoma cells from the subcutaneous tumors, the blood, and from liver metastases. The markers and gating strategies used to distinguish human melanoma cells from mouse cells are shown in Extended Data Fig. 1b. RNA sequencing revealed that interferon regulated genes were among the most significantly upregulated gene sets in melanoma cells from the blood as compared to primary subcutaneous tumors (Figure 1a). We scored each melanoma specimen for the expression of interferon regulated genes (from the Gene Ontology Biological Process “Response to type I interferon” gene set).

**Figure 1.**
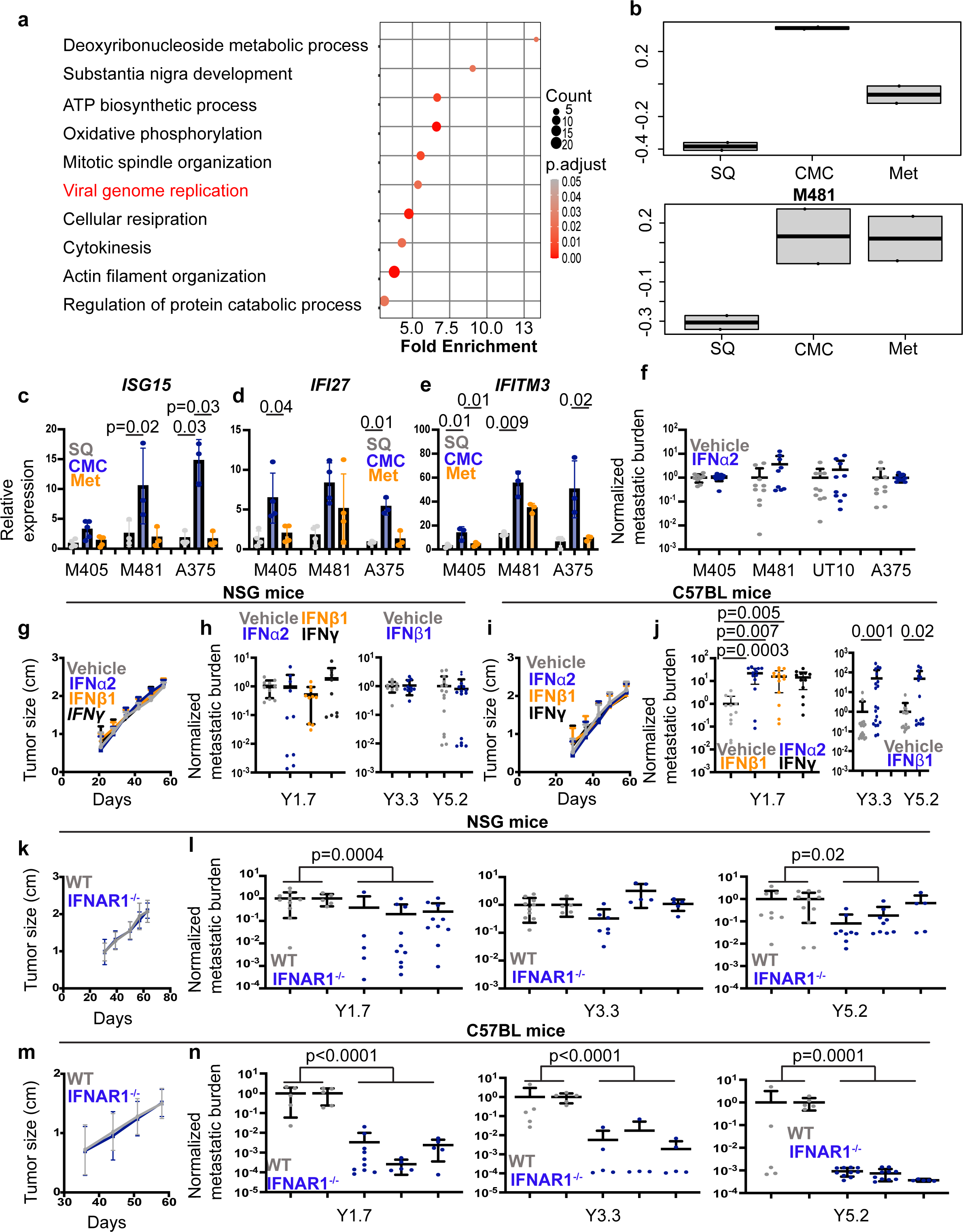
Higher interferon-regulated gene expression in metastasizing melanoma cells and increased formation of metastatic tumors after interferon treatment. a, We performed RNA sequencing on patient-derived xenograft cells (M405 and M481) isolated by flow cytometry from subcutaneous tumors, the blood, and metastatic tumors in NSG mice. After eliminating cell cycle-related genes, the 10 most significantly enriched gene sets in melanoma cells from the blood included ‘viral genome replication’ (red), which contains interferon-regulated genes. b, Interferon-regulated genes were more highly expressed by circulating (CMC) and metastatic as compared to primary subcutaneous (SQ) melanoma cells by gene set variation analysis (“Response to type I interferon” gene set). c-e, By qRT-PCR, transcript levels for the interferon-regulated genes *ISG15, IFI27* and *IFITM3* were higher in melanoma cells isolated from the blood as compared to subcutaneous or metastatic tumors of xenografted mice (two to three independent experiments per melanoma with a total of 3-5 mice per melanoma). f, Luciferase-expressing human melanoma cells were cultured overnight in 10 ng/mL hIFNa2 or vehicle and then intravenously injected into NSG mice. Metastatic disease burden was assessed five to nine weeks later by bioluminescence imaging of visceral organs and normalized to controls (two experiments per melanoma with a total of nine to ten mice per melanoma). g-j, Luciferase-expressing YUMM1.7, YUMM3.3, or YUMM5.2 mouse melanoma cells were cultured overnight in 10 ng/mL mIFN51, mIFNa2, mIFNy , or vehicle control and then injected subcutaneously (g, i) or intravenously (h, j) into NSG (g-h) or C57BL mice (i-j). We observed no differences in subcutaneous tumor growth (representative data are shown for YUMM1.7 cells; g, i). Interferon-treated mice gave rise to significantly higher tumor burden after intravenous injection into C57BL but not NSG mice (two independent experiments with a total of 9-20 mice per melanoma). k-n, *IFNAR* mutant or control mouse melanoma cells were injected subcutaneously (k, m) or intravenously (l, n) into NSG (k, l) or C57BL mice (m, n). *IFNAR* mutant and control cells did not differ in the size of the subcutaneous tumors they formed in NSG (k) or C57BL (m) mice (one experiment using YUMM1.7 cells is shown for each mouse strain, representative of 2 to 3 independent experiments for each of 3 melanomas). *IFNAR* mutant cells generally gave rise to fewer tumors in visceral organs than control cells after intravenous injection in NSG (l) and C57BL (n) mice, though the difference was much greater in C57BL mice. For each melanoma, two control clones and three independently-targeted *IFNAR1* mutant clones were studied in two to four independent experiments per melanoma with a total of 10-25 mice per melanoma. Each dot represents a different mouse and all data represent mean ± s.d. Statistical significance was assessed using repeated measures one-way (c) or two-way ANOVAs (d-e) with Dunnett’s multiple comparisons adjustments (c-e), Mann-Whitney tests followed by Holm-Sidak’s multiple comparisons adjustments (f, Y3.3 and Y5.2 of h and j, l, and n), nparLD tests (g, i, k, and m) followed by FDR multiple comparisons adjustments (g and i), or Kruskal-Wallis tests with Dunn’s multiple comparisons adjustments (for Y1.7 of h and j). All statistical tests were two-sided. No statistically significant differences were observed in f, g-i, k, or m.

Melanoma cells from the blood and metastatic tumors exhibited higher expression of interferon regulated genes as compared to melanoma cells from primary subcutaneous tumors (Figure 1b). This raised the possibility that metastasizing melanoma cells were more likely to have been exposed to interferon than other cells in primary tumors.

We assessed the levels of interferon regulated gene transcripts by quantitative RT-PCR in human melanoma cells isolated by flow cytometry from primary subcutaneous tumors, the blood, and lung metastases from NSG mice xenografted with M405 or M481 patient-derived xenografts or the A375 melanoma cell line. *Interferon Stimulated gene 15* (*ISG15)*, *Interferon-alpha Inducible protein 27* (*IFI27)*, and *Interferon-induced transmembrane protein 3* (*IFITM3)* were usually significantly more highly expressed (2.4 to 28-fold) in melanoma cells from the blood as compared to primary subcutaneous and metastatic tumors (Figure 1c-e).

To test if interferon exposure alters the metastatic potential of melanoma cells, we cultured M405, M481 and UT10 patient-derived xenograft cells or A375 cells, with or without interferon alpha 2a (IFNα2a), and then injected them intravenously in NSG mice. Five to 9 weeks later we assessed metastatic disease burden by bioluminescence imaging of visceral organs (Extended Data Fig. 1d). The vast majority of mice had tumors in visceral organs. We did not observe any statistically significant differences in disease burden among NSG mice injected with IFNα2a-treated as compared to control cells (Figure 1f). This suggested that interferon treatment did not alter the clonogenicity of melanoma cells; however, effects of IFNα2a on melanoma cell immunogenicity may not have been detected in this experiment because it was done in highly immunocompromised mice. Therefore, subsequent experiments examining the effects of interferon on metastasis were performed on Yale University Mouse Melanomas (YUMM)^30^ transplanted into syngeneic C57BL mice.

We cultured YUMM1.7 (*Braf^V600E/+^;PTEN^–/–^;Cdkn2^–/–^*), YUMM3.3 (*Braf^V600E/+^;Cdkn2^–/–^*), and YUMM5.2 (Braf^V600E/+^ p53^-/-^) melanomas^30^ with or without mouse IFNα2, IFNβ1, or IFNγ. Culture in type 1 (IFNα2 or IFNβ1) or type 2 (IFNγ) interferon did not alter the clonogenic potential of melanoma cells from any of the YUMM lines as we observed no significant difference in the ability of interferon-treated versus control cells to form spheroids in culture (Extended Data Fig. 1e) or subcutaneous tumors in NSG (Figure 1g) or C57BL (Figure 1i) mice. In NSG mice, we also observed no significant difference in the total luciferase signal (disease burden) in visceral organs after intravenous injection (Figure 1h). However, after intravenous injection into C57BL mice, we observed significantly higher disease burden (15 to 50-fold) in mice transplanted with interferon-treated as compared to control cells (Figure 1j). Similar effects were observed after culture in both type I and type II interferons. Since this difference was observed in immunocompetent, but not immunocompromised, mice and after intravenous, but not subcutaneous, transplantation, the data suggested that culture in interferon may have transiently reduced the immunogenicity of melanoma cells during metastasis.

To test if type 1 interferon signaling in vivo altered the immunogenicity of melanoma cells during metastasis, we used CRISPR to introduce stop codons into the first exon of *Interferon alpha and beta receptor subunit 1* (*IFNAR1*) from YUMM1.7, YUMM3.3, and YUMM5.2 to prevent IFNAR1 translation (Extended Data Fig. 1f). There are no additional in-frame start codons in subsequent exons. For each YUMM line, we selected two control lines transfected with Cas9 and a non-targeting scrambled sgRNA as well as 3 independently targeted mutant lines. Flow cytometric analysis showed that the mutant lines exhibited little or no IFNAR1 staining on the cell surface (Extended Data Fig. 1g-i). In non-targeted control cells, culture with IFN51 or IFNγ increased PD-L1 levels on the cell surface while in *IFNAR1* mutant cells culture with IFNγ, but not IFN51, increased PD-L1 levels (Extended Data Fig. 1j-l). The *IFNAR1* mutant cells thus exhibited a loss of IFNAR1 function.

We transplanted the non-targeted control clones and *IFNAR1* mutant clones from each YUMM line subcutaneously or intravenously into NSG and C57BL mice. The *IFNAR1*-mutant and control lines did not significantly differ in their capacity to form subcutaneous tumors in either NSG or C57BL mice (Figure 1k and 1m). However, *IFNAR1*-mutant cells formed significantly fewer tumors in visceral organs after intravenous injection (Figure 1l and 1n). The magnitude of the difference was small in NSG mice (up to 3.9-fold lower; Figure 1l) but substantial in C57BL mice (130 to 1300-fold; Figure 1n). This suggested that type 1 interferon signaling does reduce the immunogenicity of metastasizing melanoma cells in vivo. NSG mice lack T, B, and NK cells, and have macrophages that are less effective at phagocytosing human cells as compared to macrophages from C57BL mice^31,32^; therefore, it is unclear from these observations to what extent the reduction in immunogenicity was driven by a change in the interaction of metastasizing cells with adaptive versus innate immune cells.

## Heterogeneity among melanoma cells in interferon-regulated gene expression in vivo

To study the heterogeneity in interferon signaling among melanoma cells at the single cell level, we generated two types of knock-in IFN reporter cells by inserting *Green fluorescent protein* (*Gfp*) into the N-terminus of the *ISG15* gene or the *IFIT3* gene in YUMM1.7, YUMM3.3, and YUMM5.2 cells such that these cells expressed a GFP-ISG15 or a GFP-IFIT3 fusion reporter (Extended Data Fig. 2a-d). *ISG15* and *IFIT3* were among the interferon-regulated genes^33,34^ that were commonly more highly expressed in melanoma cells from the blood or metastases as compared to primary tumors (e.g. Figure 1c). In human cells, *ISG15* antagonizes interferon signaling by stabilizing USP18^35^; however, this function is absent in mouse ISG15^33^, so we would not expect any effect of the fusion protein on interferon signaling in YUMMs.

Consistent with this, GFP-ISG15 and parental control cells exhibited similar increases in PD-L1 expression when cultured in IFNβ1 (Extended Data Fig. 2e).

To test if the reporters were activated by interferon, we cultured YUMM cells expressing the GFP-ISG15 or GFP-IFIT3 reporters, with or without IFNβ1 and with or without a JAK1/2 inhibitor. JAK1/2 inhibitors block interferon receptor signaling^36^. In each of YUMM1.7, YUMM3.3, and YUMM5.2, the GFP-ISG15 and GFP-IFIT3 reporters were uniformly induced by culture in IFNβ1 and this was blocked by addition of JAK1/2 inhibitor (Figure 2a and 2b).

**Figure 2.**
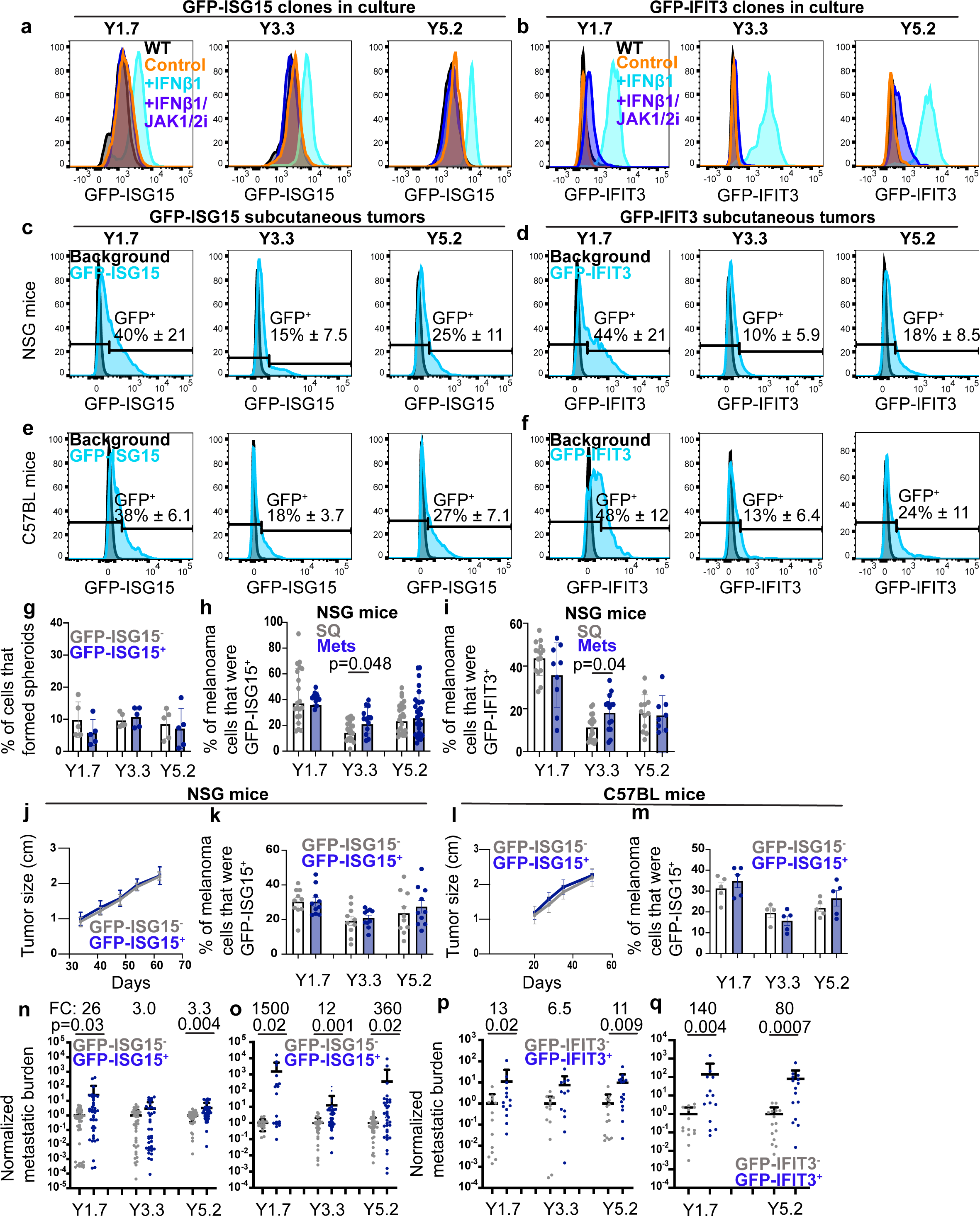
**A hyper-metastatic subset of melanoma cells**. **a-b**, Representative flow cytometry histograms depicting expression of GFP-ISG15 (**a**) or GFP-IFIT3 (**b**) fusion reporters in mouse melanoma cells after overnight culture in vehicle control or interferon 51, with or without JAK1/2 inhibitor (one representative reporter clone is shown for each melanoma). **c-f**, Representative flow cytometry histograms depicting GFP-ISG15 (**c**) or GFP-IFIT3 (**d**) reporter expression in subcutaneous mouse melanomas grown in either NSG (**c**-**d**) or C57BL mice (**e**-**f**). The percentage of reporter^+^ cells for each melanoma is shown on each plot (data represent two to five independent experiments with a total of 5-28 mice per melanoma). **g**, Percentage of GFP-ISG15^-^ or GFP-ISG15^+^ mouse melanoma cells that formed spheroids in culture after being sorted from subcutaneous tumors (two independent experiments with a total of five mice per melanoma). **h**-**i**, The percentage of melanoma cells that were GFP-ISG15^+^ (**h**) or GFP-IFIT3^+^ (**i**) in subcutaneous or metastatic tumors that formed in NSG mice after subcutaneous transplantation and spontaneous metastasis (three to five independent experiments with a total of 8-28 mice per melanoma). **j**-**m**, GFP-ISG15^-^ or GFP-ISG15^+^ cells were transplanted subcutaneously into NSG (**j, k**) or C57BL (**l, m**) mice. The rate of tumor growth was measured over time (**j, l**) as well as the percentage of GFP-ISG15^+^ melanoma cells in the subcutaneous tumors at endpoint (**k**, **m**; 2-2.5 cm in diameter; two independent experiments with a total of 4-10 mice per melanoma). **n**-**o**, GFP-ISG15^-^ or GFP-ISG15^+^ cells were transplanted intravenously into NSG (**n**) or C57BL (**o**) mice. Total metastatic tumor burden was assessed three to five weeks later by bioluminescence imaging and normalized to controls (four to seven experiments with a total of 17-42 mice per melanoma). **p, q**, A similar experiment was performed using GFP-IFIT3^-^ or GFP-IFIT3^+^ cells (two independent experiments with a total of 12-15 mice per melanoma). Each dot represents a different mouse and all data represent mean ± s.d. Statistical significance was assessed using repeated measures two-way ANOVAs followed by Sidak’s multiple comparisons adjustments (**g, l**), Student’s *t*-tests followed by Holm-Sidak’s multiple comparisons adjustments (**h-i, m,** and **p**), two-way ANOVAs with Sidak’s multiple comparisons adjustments (**k, q**), linear mixed-effects analysis (**j**), or Mann-Whitney tests with Holm-Sidak’s multiple comparison’s adjustments (**n**-**o**). All statistical tests were two-sided. No statistically significant differences were observed in **g** or **j-m**.

To assess heterogeneity among melanoma cells in type 1 interferon signaling in vivo, we transplanted GFP-ISG15 and GFP-IFIT3 YUMM cells subcutaneously into NSG and C57BL mice. The GFP-ISG15 and GFP-IFIT3 reporters were expressed in 14 to 48% of melanoma cells in primary subcutaneous tumors, depending on the YUMM line, but no significant differences were observed between the same YUMM lines in NSG versus C57BL mice (Figure 2c-f). This suggested that the interferon that drives reporter expression in vivo may come primarily from innate immune cells that exist in both NSG and C57BL mice. We allowed the melanomas to spontaneously metastasize in NSG mice and then assessed the percentage of reporter+ cells in the metastatic tumors. YUMM1.7 and YUMM5.2 exhibited similar percentages of GFP-ISG15^+^ and GFP-IFIT3^+^ cells in primary subcutaneous as compared to metastatic tumors while YUMM3.3 had significantly higher percentages of GFP-ISG15^+^ and GFP-IFIT3^+^ cells in metastatic tumors as compared to subcutaneous tumors (Figure 2h and 2i).

We isolated GFP-ISG15^+^ and GFP-ISG15^-^ cells by flow cytometry from primary subcutaneous tumors in C57BL mice and performed RNA sequencing. All 8 of the gene sets that were most significantly enriched in GFP-ISG15^+^ as compared to GFP-ISG15^-^ cells contained interferon-regulated genes (Extended Data Fig. 2f). Moreover, the ten genes that were most significantly upregulated in GFP-ISG15^+^ as compared to GFP-ISG15^-^ cells from each YUMM line were almost all interferon-regulated (Extended Data Fig. 2g-i). Interferon-regulated genes were thus broadly increased in expression within GFP-ISG15^+^ as compared to GFP-ISG15^-^ cells from each YUMM line.

To compare the functions of reporter positive and negative cells, we isolated GFP-ISG15^+^ and GFP-ISG15^-^ cells by flow cytometry. The GFP-ISG15^+^ and GFP-ISG15^-^ cells did not significantly differ in their capacity to form spheroids in culture (Figure 2g) or primary tumors after subcutaneous injection into NSG mice (Figure 2j) or C57BL mice (Figure 2l). Exposure to interferon, thus, did not alter the clonogenic potential of melanoma cells in culture or in vivo.

Moreover, the frequency of reporter positive cells in these subcutaneous tumors did not significantly differ among tumors formed by GFP-ISG15^+^ as compared to GFP-ISG15^-^ cells (Figure 2k and 2m). This demonstrates that ISG15 is not constitutively activated in YUMM cells and that GFP-ISG15^+^ cells give rise to GFP-ISG15^-^ cells, and vice versa, in vivo.

We performed additional experiments to assess whether the interferon that promoted GFP-ISG15 expression was produced autonomously by melanoma cells or non-cell-autonomously by other cells. First, we sorted unfractionated cells from GFP-ISG15-bearing melanoma cells into culture and assessed the percentage of their progeny that were GFP-ISG15^+^. In all cases, the cells gave rise exclusively to GFP-ISG15^-^ cells in culture (Extended Data Fig. 2j), suggesting that the melanoma cells did not produce enough interferon to activate reporter expression in culture and that reporter expression in tumors in vivo likely reflected non-cell-autonomous interferon production by immune cells. Consistent with this, loss of *IFNAR1* function (Extended Data Fig. 1f) significantly reduced the percentage of GFP-ISG15^+^ cells in tumors formed by every YUMM line, irrespective of whether the cells were injected into NSG or C57BL mice (Extended Data Fig. 2k-l). Furthermore, treatment of tumor-bearing mice with reverse transcriptase inhibitors or STING antagonist did not significantly affect the percentage of GFP-ISG15^+^ cells within tumors formed by any YUMM line (Extended Data Fig. 2m-o), suggesting that the interferon that drove GFP-ISG15 expression was not produced as a result of STING activation or retrotransposon expression in melanoma cells.

## A hyper-metastatic subset of melanoma cells

To assess the effect of interferon signaling on the metastatic potential of melanoma cells, we sorted GFP^+^ or GFP^-^ cells from subcutaneous tumors formed by YUMM1.7, YUMM3.3, or YUMM5.2 melanomas expressing the GFP-ISG15 or GFP-IFIT3 reporters and then injected the cells intravenously into NSG or C57BL mice. In NSG mice, the GFP^+^ cells marked with either reporter usually formed significantly more tumors (3.0 to 26-fold increase in disease burden within visceral organs) than the GFP^-^ cells after intravenous injection (Figure 2n and 2p). In C57BL mice, the GFP^+^ cells marked with either reporter also formed significantly more tumors than the GFP^-^ cells after intravenous injection but the magnitude of the difference was much larger: 12 to 1500-fold increase in disease burden within visceral organs (Figure 2o and 2q).

This continued to suggest that melanoma cells that expressed the GFP-ISG15 or GFP-IFIT3 reporters in vivo were far better at forming metastatic tumors, particularly in an immunocompetent environment.

The interferon-regulated genes that were significantly more highly expressed in GFP-ISG15^+^ cells as compared to GFP-ISG15^-^ cells included several that are known to regulate immunogenicity^13^, including *beta-2-microglobulin (B2M), CD274 (PD-L1), major histocompatibility complex type 1* (*H2kb*), and *CD47* (Extended Data Fig. 2f-i). We confirmed by flow cytometry that protein levels for each of these gene products increased as GFP-ISG15 expression increased (Figure 3a and 3b).

**Figure 3.**
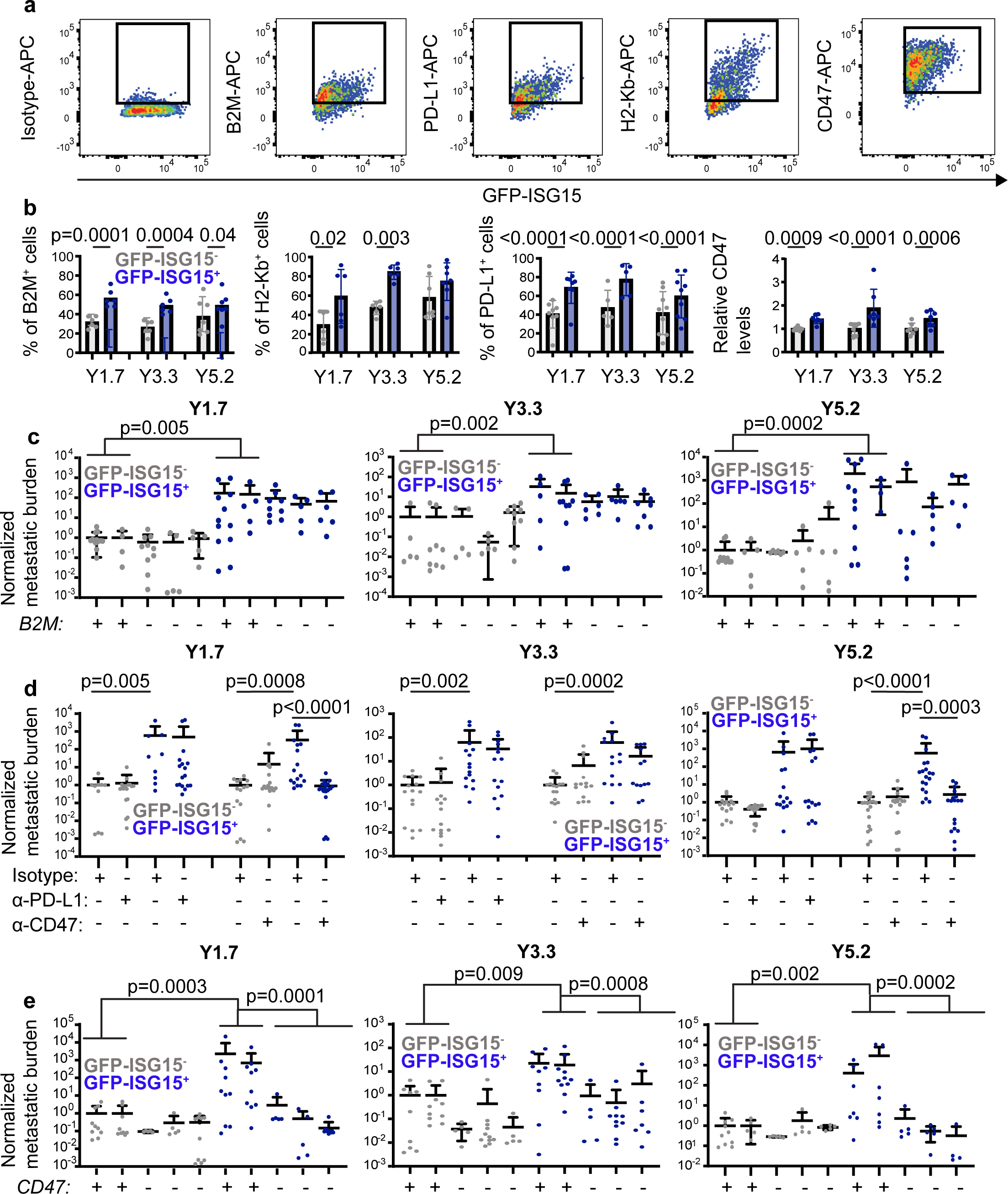
**Increased CD47 levels contribute to the increased metastatic potential of GFP-ISG15^+^ cells. a-b**, Representative flow cytometry plots (**a**) and quantification (**b**) of beta-2-microglobulin (B2M), PD-L1, H2-Kb, and CD47 levels on the surface of GFP-ISG15^-^ versus GFP-ISG15^+^ melanoma cells from subcutaneous tumors (two independent experiments with a total of 5-9 mice per melanoma). **c**, C57BL mice were injected intravenously with GFP-ISG15^-^ or GFP-ISG15^+^ cells from *B2M* mutant (-) or control (+) melanomas and metastatic disease burden was assessed three to five weeks later by bioluminescence imaging of visceral organs. For each melanoma (YUMM1.7, 3.3, or 5.2), two control clones and three independently-targeted *B2M* mutant clones were studied in two to three independent experiments with a total of 14-20 mice per melanoma. **d**, GFP-ISG15^-^ or GFP-ISG15^+^ cells were treated with blocking antibodies against PD-L1 or CD47 or isotype control antibody then injected intravenously into C57BL mice and metastatic disease burden was assessed three to five weeks later by bioluminescence imaging (two to three experiments with a total of 9-21 mice per melanoma). **e**, C57BL mice were injected intravenously with GFP-ISG15^-^ or GFP-ISG15^+^ cells from *CD47* mutant (-) or control (+) melanomas then metastatic disease burden was assessed three to five weeks later. For each melanoma, two control clones and three independently-targeted *CD47* mutant clones were studied in two to four independent experiments with a total of 14-22 mice per melanoma. Each dot represents cells from a different mouse and all data represent mean ± s.d. Statistical significance was assessed using repeated measures two-way ANOVAs with Sidak’s multiple comparisons adjustments (**b**), Kruskal-Wallis tests with Dunn’s multiple comparisons adjustments (**c**, Y1.7 and Y5.2 of **d**, and **-e**), or a one-way ANOVA with Sidak’s multiple comparisons adjustment (Y3.3 of **d**). All statistical tests were two-sided.

To test whether any of these gene products mediated the increased capacity of GFP-ISG15^+^ cells as compared to GFP-ISG15^-^ cells to form tumors after intravenous injection, we first used CRISPR to introduce stop codons into the first exon of *beta-2-microglobulin (B2M*) in YUMM1.7, YUMM3.3, and YUMM5.2 melanomas to prevent translation (Extended Data Fig. 3a). B2M is a class I major histocompatibility complex (MHC) component that is necessary for surface expression of class I MHC (including H2-Kb)^37^. For each YUMM line, we selected two control lines transfected with Cas9 and a non-targeting scrambled sgRNA as well as 3 independently targeted mutant lines. Flow cytometric analysis showed that none of the mutant lines exhibited detectable B2M or H2-Kb staining on the cell surface (Extended Data Fig. 3b and 3c). We intravenously injected GFP-ISG15^-^ cells or GFP-ISG15^+^ cells from the *B2M* mutant and control melanomas into C57BL mice and then quantitated metastatic disease burden in visceral organs four to five weeks later. We did not observe any significant differences in metastatic disease burden between mice injected with *B2M* mutant as compared to control melanoma cells, irrespective of whether we injected GFP-ISG15^-^ cells or GFP-ISG15^+^ cells (Figure 3c).

This suggested that the increased B2M expression by GFP-ISG15^+^ cells was not responsible for the increased capacity of these cells to form metastatic tumors in immunocompetent mice.

We next tested if the increased metastatic potential of GFP-ISG15^+^ cells was driven by increased PD-L1 or CD47 on the cell surface. We treated GFP-ISG15^+^ cells or GFP-ISG15^-^cells from YUMM1.7, YUMM3.3, and YUMM5.2 melanomas with blocking antibodies against PD-L1, CD47, or a rat IgG2b isotype control antibody and then intravenously injected the cells into C57BL mice. Four to five weeks later we analyzed metastatic disease burden in visceral organs by bioluminescence imaging. Among mice injected with cells treated with isotype control antibody, mice injected with GFP-ISG15^+^ cells had significantly greater metastatic disease burdens as compared to mice injected with GFP-ISG15^-^ cells (Figure 3d). Among mice injected with cells treated with anti-PD-L1 antibody, we observed no significant difference in metastatic disease burden between mice transplanted with anti-PD-L1 versus isotype control antibody treated cells for any of the YUMM lines, irrespective of whether the mice were injected with GFP-ISG15^+^ cells or GFP-ISG15^-^ cells (Figure 3d). This suggested that the increased PD-L1 expression by GFP-ISG15^+^ cells was not responsible for the increased capacity of these cells to form metastatic tumors. We also observed no significant difference in metastatic disease burden between mice transplanted with GFP-ISG15^-^ cells that were treated with anti-CD47 versus isotype control antibody; however, we observed significantly reduced metastatic disease burden in mice transplanted with GFP-ISG15^+^ cells that were treated with anti-CD47 as compared to isotype control antibody for YUMM1.7 (220-fold reduction) and YUMM5.2 (125-fold reduction) (Figure 3d). Anti-CD47 antibody treatment was associated with a trend toward reduced tumor formation in YUMM3.3 cells (3.8-fold reduction) but the difference was not statistically significant (Figure 3d). This suggested CD47 contributed to the increased ability of GFP-ISG15^+^ cells to form metastatic tumors as compared to GFP-ISG15^-^ cells at least in YUMM1.7 and YUMM5.2.

To further explore this through an orthogonal approach, we used CRISPR to introduce stop codons into the first exon of *CD47* in YUMM1.7, YUMM3.3, and YUMM5.2 melanomas to prevent translation (Extended Data Fig. 3d). For each YUMM line, we selected two control lines transfected with Cas9 and a non-targeting scrambled sgRNA as well as 3 independently targeted mutant lines. Flow cytometric analysis showed that none of the mutant lines exhibited detectable CD47 staining on the cell surface (Extended Data Fig. 3e). We intravenously injected GFP-ISG15^-^ cells or GFP-ISG15^+^ cells from the *CD47* mutant and control melanomas into C57BL mice and then quantitated metastatic disease burden in visceral organs four to five weeks later. Among mice injected with GFP-ISG15^-^ cells, *CD47* mutant and control cells did not significantly differ in the disease burden they gave rise to (Figure 3e). Conversely, among C57BL mice injected with GFP-ISG15^+^ cells, the *CD47* mutant cells always gave rise to significantly reduced disease burden for all three YUMMs (15 to 1600-fold reductions; Figure 3e). This further demonstrated that CD47 contributed to the increased ability of GFP-ISG15^+^ cells to form metastatic tumors as compared to GFP-ISG15^-^ cells in immunocompetent mice.

### GFP-ISG15^+^ melanoma cells are resistant to phagocytosis

Since CD47 inhibits phagocytosis by macrophages and other myeloid cells^38–41^, we tested whether GFP-ISG15^+^ cells are more resistant than GFP-ISG15^-^ cells to phagocytosis by macrophages in culture. We tested this by isolating GFP-ISG15^+^ cells and GFP-ISG15^-^ cells from YUMM1.7, YUMM3.3, and YUMM5.2 subcutaneous tumors, labelling them with carboxyfluorescein succinimidyl ester (CFSE) dye, co-culturing with C57BL macrophages overnight, and then quantitating the percentage of CFSE^+^ macrophages by flow cytometry. We observed significantly reduced phagocytosis of GFP-ISG15^+^ cells as compared to GFP-ISG15^-^cells (Figure 4a and 4b). To assess whether CD47 contributed to this difference, we performed the same experiment on GFP-ISG15^+^ cells and GFP-ISG15^-^ cells from CD47 mutant melanomas. We observed significantly more phagocytosis of CD47 mutant as compared to control melanoma cells (Figure 4a and 4b). In the absence of CD47, the phagocytosis of GFP-ISG15^+^ cells and GFP-ISG15^-^ cells did not significantly differ in YUMM1.7 and YUMM3.3, though there was still reduced phagocytosis of GFP-ISG15 ^+^ cells as compared to GFP-ISG15^-^cells in YUMM5.2 (Figure 4b). This demonstrated that GFP-ISG15^+^ cells were more resistant to phagocytosis by macrophages than GFP-ISG15^-^ cells and that CD47 contributed to this difference, but that other mechanisms also contributed, particularly in YUMM5.2 cells.

**Figure 4.**
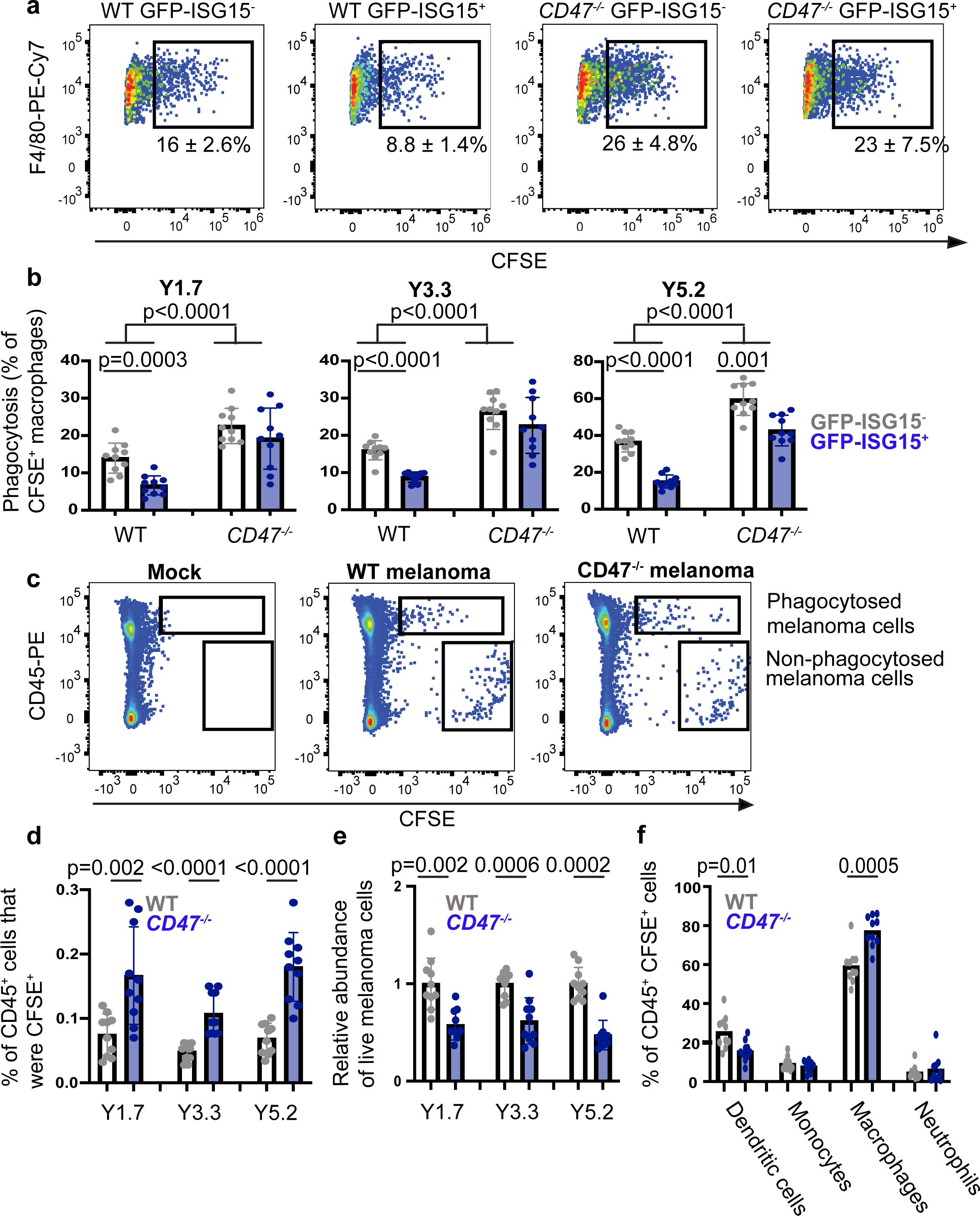
**CD47 protects ISG15^+^ melanoma cells from macrophage-mediated phagocytosis. a-b**, CFSE-labeled mouse melanoma cells (YUMM1.7, 3.3, or 5.2) were co-cultured with C57BL bone marrow-derived macrophages, then the percentage of F4/80^+^ macrophages that had phagocytosed a melanoma cell (CFSE^+^) was determined by flow cytometry 16-20 hours later. The melanoma cells were GFP-ISG15^-^ or GFP-ISG15^+^ and were obtained from *CD47* mutant or control melanomas (two experiments with a total of 10 mice per melanoma). **c**-**d**, C57BL mice were injected intravenously with *CD47* mutant or control melanoma cells from subcutaneous tumors. The next day, lungs were harvested, dissociated and the percentage of CD45^+^ cells that were CFSE^+^ was determined by flow cytometry (two experiments per melanoma with a total of 10 mice per melanoma). **e**, Mice in panels **c, d** transplanted with *CD47* mutant melanoma cells had fewer non-phagocytosed melanoma cells per lung as compared to mice transplanted with control cells (two experiments with 10 mice per melanoma). **f**, The percentage of CD45^+^CFSE^+^ lung cells in panels **c, d** that were dendritic cells, monocytes, macrophages or neutrophils (two experiments with ten mice per melanoma). Each dot represents a different mouse and all data represent mean ± s.d. Statistical significance was assessed using repeated measures two-way ANOVAs followed by Sidak’s multiple comparisons adjustments (**b**), or Student’s *t*-tests (**d-f**) or Mann-Whitney tests (**e**) followed by Holm-Sidak’s multiple comparisons adjustments (**d-f**). All statistical tests were two-sided.

We performed a similar experiment in vivo by intravenously injecting CFSE-labelled melanoma cells from CD47 mutant or control subcutaneous tumors into C57BL mice. The next day, we harvested the lungs, enzymatically dissociated, and assessed the frequency of CD45^+^CFSE^+^ cells by flow cytometry (Figure 4c). The mice injected with CD47 mutant melanoma cells had significantly higher percentages of CD45^+^ cells that were CFSE^+^ as compared to mice injected with control cells (Figure 4c and 4d). Consistent with this, the numbers of melanoma cells remaining in the lungs of mice injected with CD47 mutant cells were significantly lower than in mice injected with control cells (Figure 4e). Consistent with a prior study^42^, most of the CD45^+^CFSE^+^ cells in the lung were macrophages, along with smaller numbers of dendritic cells and rare monocytes and neutrophils (Figure 4f). CD47 deficiency significantly increased the frequency of CFSE^+^ macrophages in the lung (Figure 4f). CD47 thus protected melanoma cells from phagocytosis by macrophages in the lung.

### GFP-ISG15^+^ cells co-localize with macrophages in inflamed tumor regions

Since ISG15 expression is driven by type 1 interferon, we assessed *Ifna* and *Ifnb* expression among cells within primary subcutaneous YUMMs. We isolated GFP-ISG15^+^ melanoma cells, GFP-ISG15^-^ melanoma cells, F4/80^+^CD11b^+^CD24^-^CD45^+^ macrophages, CD11b^+^Ly6C^+^Ly6G^-^CD88^+^CD11c^-^CD45^+^ monocytes, CD24^+^CD11c^+^CD45^+^ dendritic cells, CD3^+^CD11b^-^B220^-^CD45^+^ T cells, B220^+^CD3^+^CD11b^-^CD45^+^ B cells, and NK1.1^+^CD11b^+^F4/80^-^CD45^+^ natural killer (NK) cells by flow cytometry and then performed quantitative RT-PCR for *Ifna* and *Ifnb* (see Extended Data Fig. 4a for the flow cytometry gates used to isolate these cells). Macrophages and monocytes expressed the highest levels of *Ifna* and *Ifnb*, followed by GFP-ISG15^+^ melanoma cells (Figure 5a and 5b).

**Figure 5.**
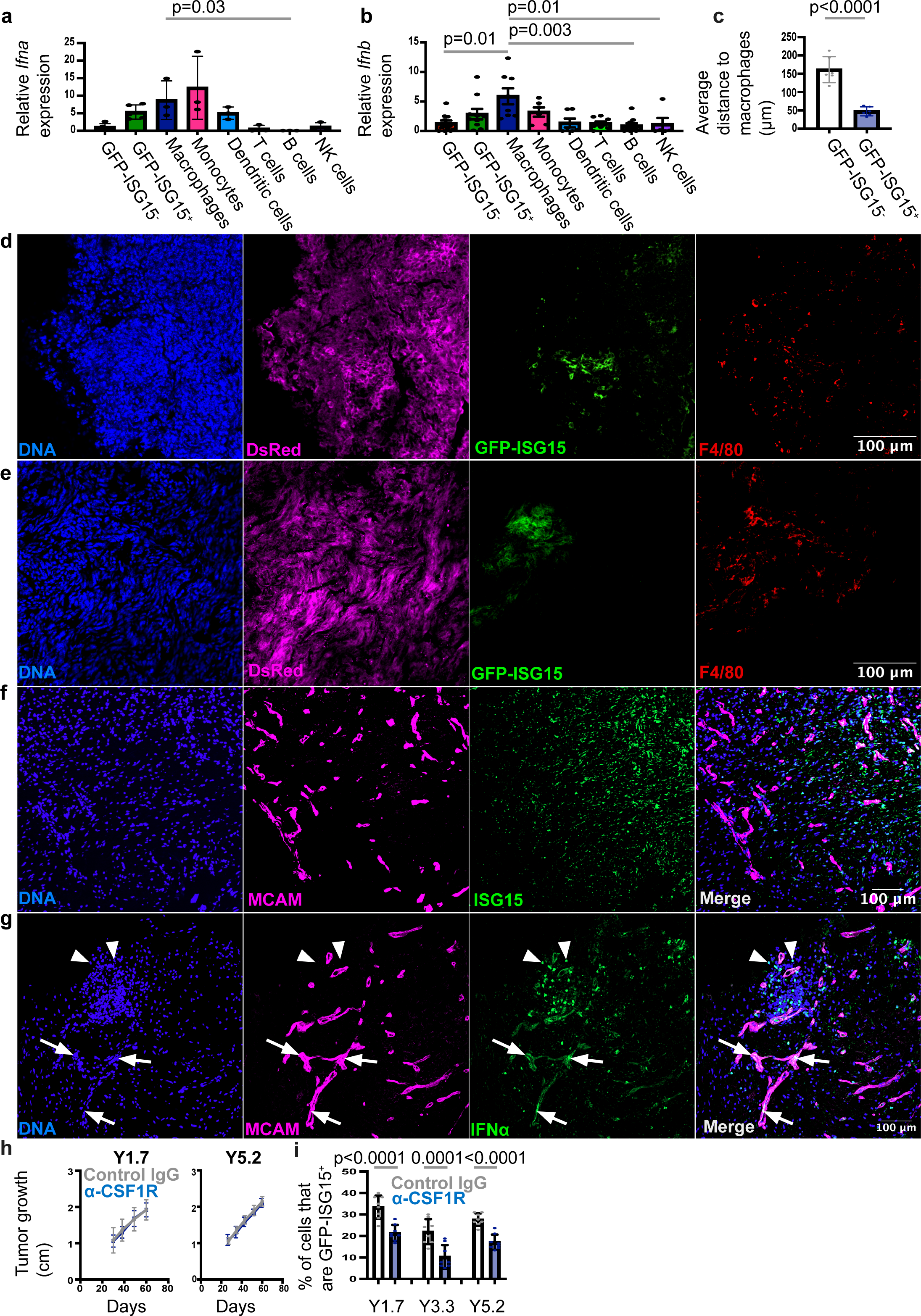
GFP-ISG15^+^ melanoma cells co-localize with macrophages within inflamed regions of primary tumors. a, **b**, qRT-PCR analysis of *Ifna* (**a**) and *Ifnb* (**b**) transcript levels in sorted GFP-ISG15^-^ or GFP-ISG15^+^ mouse melanoma cells or tumor-infiltrating immune cells (two to three independent experiments with a total of 3 to 10 melanomas from different mice). The markers and flow cytometry gates used to sort immune cells are shown in Extended Data Fig. 4a (CD115 was used to sort monocytes from the lungs while CD88 was used to sort tumor-associated monocytes). **c**, The average distance from GFP-ISG15^-^ or GFP-ISG15^+^ melanoma cells to macrophages within mouse melanoma specimens (six independent experiments with a total of six melanomas from different mice). **d**, **e**, Sections from subcutaneous tumors formed by YUMM1.7 GFP-ISG15^+^ melanoma cells in C57BL mice were stained with DAPI (blue) to label nuclei, anti-DsRed antibody (fuchsia) to label melanoma cells, anti-GFP antibody (green) to label GFP-ISG15^+^ cells, and anti-F4/80 antibody (red) to label macrophages. These tumors were dissected from mice before sectioning to eliminate surrounding normal tissue. **f, g**, Melanoma specimens that were surgically resected from humans were stained with DAPI (blue), anti-MCAM antibody (fuchsia) to label melanoma cells and anti-ISG15 (**c**, green) or anti-IFNα (**d**, green) antibody to label interferon-stimulated or interferon-producing cells, respectively. **h**, **i** Subcutaneous tumor growth (**h**) in C57BL mice injected subcutaneously with melanoma cells. Once tumors were palpable, mice were injected twice weekly with anti-CSF1R or isotype control antibody. When tumors reached 2-2.5 cm in diameter, the percentage of melanoma cells that were GFP-ISG15^+^ was assessed by flow cytometry (**i**, two experiments with a total of 5-8 mice per melanoma). Each dot represents a different mouse/melanoma and all data represent mean ± s.d. Statistical significance was assessed using Kruskal-Wallis tests followed by Dunn’s multiple comparisons adjustments (**a, b**), a Student’s *t*-test (**c**), nparLD tests (**h**), or a two-way ANOVA followed by Sidak’s multiple comparisons adjustment (**i**). All statistical tests were two-sided. No statistically significant differences were observed in (**h**).

In melanomas^2,24,26,27^ and other cancers^23,25,43^, T cells, other immune cells, and inflammatory cytokines cluster within certain regions of tumors. To assess the localization of GFP-ISG15^+^ cells, we performed immunofluorescence analysis on sections from primary mouse and human melanomas. We first examined sections from subcutaneous tumors formed by GFP-ISG15 YUMMs in C57BL mice. GFP-ISG15-expression was not uniform throughout the tumors. Rather GFP-ISG15^+^ cells clustered in tumor regions that were also enriched for F4/80^+^ macrophages (Figure 5d and 5e). Consistent with this, GFP-ISG15^+^ cells were much closer than GFP-ISG15^-^ cells, on average, to macrophages (Figure 5c). ISG15^+^ cells also clustered in certain regions of primary cutaneous melanomas obtained directly from patients (Figure 5f).

These tumors also exhibited clusters of IFNα^+^ cells, which included IFNα^+^ melanoma cells (arrows in Figure 5g) and IFNα^+^ non-melanoma cells (arrowheads in Figure 5g). The IFNα^+^ non-melanoma cells included CD68^+^ monocytes/macrophages (Extended Data Fig. 4d).

To test if monocytes/macrophages were a functionally important source of type 1 IFN for interferon-regulated gene activation in melanoma cells, we depleted them by treating melanoma-bearing mice with anti-Colony Stimulating Factor 1 Receptor (CSF1R) or isotype control antibody^44^, beginning when subcutaneous tumors became palpable. Treatment with anti-CSF1R nearly completely eliminated macrophages as well as depleting monocytes from primary subcutaneous melanomas (Extended Data Fig. 4c). Anti-CSF1R antibody did not significantly affect primary tumor growth (Figure 5h) but did significantly reduce the percentage of melanoma cells in primary tumors that were GFP-ISG15^+^ (Figure 5i). Macrophages/monocytes are thus a significant source of interferon for the activation of *ISG15* expression in melanoma cells but they are not the only source. Type 1 interferon produced by GFP-ISG15^+^ melanoma cells themselves (Figure 5a and 5b) may contribute to the expression of interferon-regulated genes.

## Macrophages are a double-edged sword

In melanoma, macrophages and monocytes inhibit the development of metastatic disease in some contexts^45,46^ while promoting the development of metastatic disease in other contexts^47–50^. To test if macrophage/monocyte depletion reduced the metastatic potential of melanoma cells from primary tumors, we isolated GFP-ISG15^+^ cells and GFP-ISG15^-^ cells by flow cytometry from subcutaneous tumors growing in C57BL mice treated with anti-CSF1R or isotype control antibody. Anti-CSF1R antibody treatment did not significantly affect the ability of GFP-ISG15^-^ cells to give rise to metastatic disease after intravenous injection into C57BL mice when compared to isotype control antibody treatment (Figure 6a). In contrast, anti-CSF1R antibody treatment did significantly reduce (11 to 46-fold) the ability of GFP-ISG15^+^ cells to give rise to metastatic disease after intravenous injection into C57BL mice (Figure 6a). This suggests that in our experiments, macrophages/monocytes increased the metastatic potential of ISG15^+^ but not ISG15^-^ melanoma cells. The context-dependence of the effects of macrophages on metastatic potential can thus be resolved partly by distinguishing between effects on melanoma cells that differ in terms of interferon-regulated gene expression.

**Figure 6.**
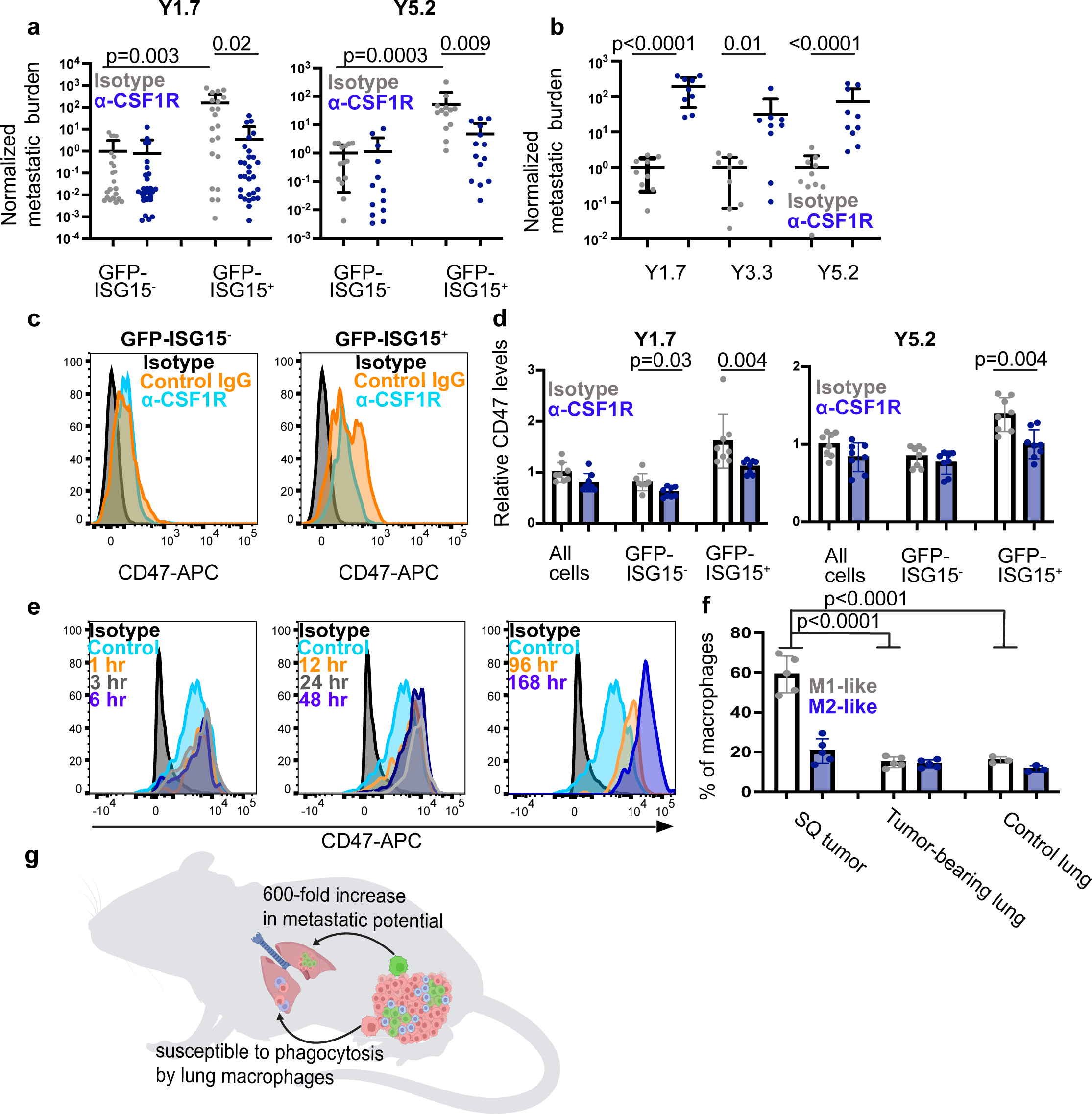
Macrophages are a double-edged sword: macrophage depletion in donor mice reduced the metastatic potential of GFP-ISG15^+^ melanoma cells but macrophage depletion in recipient mice increased the formation of metastatic tumors after transplantation of melanoma cells from untreated donor mice. **a**, GFP-ISG15^-^ or GFP-ISG15^+^ melanoma cells sorted from primary tumors in anti-CSF1R or isotype control antibody-treated donor mice were injected intravenously into C57BL recipient mice to assess the effect of macrophage/monocyte depletion in donor mice on metastatic potential. Metastatic disease burden was assessed three to five weeks later by bioluminescence imaging and normalized to control mice (three to four experiments with a total of 13-29 mice per melanoma). **b**, C57BL recipient mice were treated with anti-CSF1R or isotype control antibody for two weeks to deplete macrophages/monocytes and then injected intravenously with unfractionated melanoma cells from untreated donor mice to assess the effect of systemic macrophage/monocyte depletion in recipient mice on the ability of melanoma cells from untreated donor mice to form metastatic tumors (two experiments with a total of 9-10 mice per melanoma). **c-d**, CD47 levels in GFP-ISG15^-^ or GFP-ISG15^+^ melanoma cells from primary tumors in C57BL mice treated with either anti-CSF1R or isotype control antibody (one histogram per treatment, representative of data from two experiments with a total of eight mice per melanoma). **e**, Representative histograms of CD47 staining in mouse melanoma cells that were cultured in 10 ng/ml interferon 51 or vehicle (control) for varying amounts of time. **f,** The percentages of F4/80^+^Ly6c^-^macrophages that were M1-like (CD86^+^CD206^-^CD163^-^ cells) or M2-like (CD163^+^CD206^+^CD86^-^cells) in primary subcutaneous melanomas, lungs containing micrometastases, and lungs from non-injected control C57BL mice (one experiment with a total of 3-5 mice). **g**, Macrophages/monocytes in established primary tumors contribute to the creation of a hyper-metastatic subset of melanoma cells that can be distinguished based on the expression of interferon-regulated genes, including CD47, which protects these cells from phagocytosis by macrophages at sites of metastasis. Each dot represents a different mouse and all data represent mean ± s.d. Statistical significance was assessed by a Kruskal-Wallis test followed by Dunn’s multiple comparisons adjustment (**a**), Student’s *t*-tests or Mann-Whitney tests followed by Holm-Sidak’s multiple comparisons adjustments (**b**), or one-way ANOVAs followed by Sidak’s (**d**) or Tukey’s multiple comparisons adjustment (**f**). All statistical tests were two-sided.

We further hypothesized that macrophages could also have opposing effects in primary tumors as compared to sites of metastasis, where macrophages phagocytose disseminated cancer cells^45,46^. To test this, we depleted macrophages in normal C57BL mice by treating them with anti-CSF1R or isotype control antibody and then intravenously injected unfractionated YUMM1.7, YUMM3.3, or YUMM5.2 melanoma cells obtained from subcutaneous tumors. Each melanoma gave rise to significantly higher (31 to 190-fold) metastatic disease burden in mice treated with anti-CSF1R as compared to isotype control antibody (Figure 6b). This confirms that macrophages represent an important defense against disseminated cancer cells, substantially reducing the percentage of cells that survived to form metastatic tumors. Macrophages are thus a double-edged sword, increasing the metastatic potential of a subset of melanoma cells in established primary tumors while eliminating disseminated cancer cells from sites of metastasis.

One important difference between melanoma cells in established primary tumors as compared to sites of distant metastasis is that melanoma cells in inflamed regions of established primary tumors are persistently exposed to interferon while disseminated melanoma cells are unlikely to be (prior to the formation of metastatic tumors). ISG15^+^ melanoma cells had dramatically increased metastatic potential as compared to ISG15^-^ melanoma cells (Figure 2o) partly due to increased CD47 expression (Figure 3d and 3e), which inhibits phagocytosis by macrophages at metastatic sites (Figure 4c-f). This raised the question of whether CD47 expression by melanoma cells in primary tumors was influenced by macrophages/monocytes or the duration of exposure to interferon. To test this, we treated melanoma-bearing C57BL mice with anti-CSF1R or isotype control antibody and then assessed CD47 levels on the surface of GFP-ISG15^+^ and GFP-ISG15^-^ cells by flow cytometry. Anti-CSF1R antibody did not significantly affect CD47 expression by GFP-ISG15^-^ cells but did significantly reduce CD47 expression by GFP-ISG15^+^ cells (Figure 6c and 6d). Moreover, culture of melanoma cells in IFN-β1 for 1 to 6 hours had little effect on CD47 levels but CD47 levels began to increase after culture in IFN-β1 for 12 to 48 hours and continued to increase through 4 to 7 days (Figure 6e). This suggests that the prolonged exposure of a subset of melanoma cells to interferon in established primary tumors, partly produced by macrophages/monocytes, increased the metastatic potential of these cells, partly by increasing CD47 expression.

Most F4/80^+^Ly6C^-^ macrophages in primary subcutaneous tumors were M1-like (CD86^+^CD206^-^CD163^-^ cells)^51,52^ while most macrophages in the lungs of tumor-bearing or non-tumor bearing mice were not (Figure 6f). Approximately 20% of F4/80^+^Ly6c^-^ macrophages in primary subcutaneous tumors and lung were M2-like (CD163^+^CD206^+^CD86^-^ cells)^51,52^ (Figure 6f). This suggests there were differences in the composition of the macrophage pool in established primary tumors as compared to sites of metastasis. Overall, the data indicate that macrophages/monocytes in established primary tumors contribute to the creation of a hyper-metastatic subset of melanoma cells that can be distinguished based on the expression of interferon-regulated genes, including CD47, which protects these cells from phagocytosis at sites of metastasis (Figure 6g).

## Discussion

Myeloid cells and inflammation have context-dependent effects on cancer progression, sometimes promoting and sometimes inhibiting^51,53–55^. Our study suggests that that the context dependence of these effects, even within the same tumor, is driven partly by changes in cancer cells as a consequence of interferon signaling. Cells in inflamed regions of established primary tumors are likely persistently exposed to interferon over a period of days or weeks, leading to the development of a hyper-metastatic subset of cells that evades phagocytosis after dissemination. In contrast, recently disseminated cancer cells are not likely to be persistently exposed to interferon in sites of nascent metastasis, leaving them unable to acquire phagocytosis resistance after dissemination. Macrophages/monocytes thus inhibit cancer progression by eliminating scattered, disseminated cancer cells at sites of metastasis but their infiltration into inflamed regions of established primary tumors appears to promote cancer progression by persistently activating interferon signaling in a subset of melanoma cells, increasing their metastatic potential.

Macrophages and monocytes are an important source of interferon that increases the metastatic potential of a subset of primary melanoma cells (Figure 5). Macrophages have also been shown to promote metastasis through other mechanisms, including stimulating cancer cell motility/invasion^56,57^, establishing premetastatic niches that facilitate the survival of disseminated cancer cells^48,49,58^ and restraining T cell activity^59^. Thus, macrophages promote cancer progression through multiple mechanisms.

Interferon has been used as a stand-alone therapy or adjuvant therapy for melanoma, but this did not increase overall survival^60^. Trials that combined interferon with chemotherapy also did not improve overall survival^61^. JAK1/2 inhibitors, which inhibit interferon signaling, synergize with immune checkpoint blockade in melanoma and other cancers by altering the functions of myeloid cells and T cells in tumors^14,36,62,63^. Our results raise the possibility that JAK inhibitors might also act directly on cancer cells to reduce the expression of CD47 by cancer cells stimulated by persistent interferon. This would be consistent with the clinical observation that CD47 inhibition can synergize with immune checkpoint blockade by increasing cancer cell phagocytosis and antigen presentation for T cell activation^64–67^.

## ACKNOWLEDGEMENTS

S.J.M. is a Howard Hughes Medical Institute (HHMI) Investigator, the Mary McDermott Cook Chair in Pediatric Genetics, the Kathryn and Gene Bishop Distinguished Chair in Pediatric Research, the director of the Hamon Laboratory for Stem Cells and Cancer, and a Cancer Prevention and Research Institute of Texas Scholar. The research was supported by the Cancer Prevention and Research Institute of Texas (RP170114 and RP180778) and the National Institutes of Health (U01 CA228608 and U54 CA268072). We thank N. Meireles (University of Michigan) for collecting human melanomas, the BioHPC (High Performance Computing) core for computational resources and data storage, M. Nitcher and D.C. for mouse colony management, M. Ortiz and the Moody Foundation Flow Cytometry Facility, Y. Kim and the Children’s Research Institute Sequencing Facility, the University of Texas Southwestern Medical Center Tissue Management Shared Resource, a shared resource of the Simmons Comprehensive Cancer Center, which is supported partly by the National Cancer Institute (P30 CA142543) as well as M. Mettlen and the Quantitative Light Microscopy Core Facility. This article is subject to HHMI’s Open Access to Publications policy. HHMI lab heads have previously granted a nonexclusive CC BY 4.0 license to the public and a sublicensable license to HHMI in their research articles. Pursuant to those licenses, the author-accepted manuscript of this article can be made freely available under a CC BY 4.0 license immediately upon publication.

## AUTHOR CONTRIBUTIONS

M.H.M. and S.J.M. conceived the project, designed, and interpreted experiments. M.H.M performed most of the experiments. D.C. transplanted melanoma cells into mice and performed bioluminescence imaging in some experiments. T.W. helped to perform CRISPR editing of mouse melanomas. T.W. and D.L.M. assisted with flow cytometry. A.S. obtained human melanoma specimens. E.P. performed RNA sequencing on human melanoma cells from NSG mice. Z.Z. and B.K. performed statistical analyses. M.H.M. and S.J.M. wrote the manuscript.

## AUTHOR INFORMATION

Correspondence and requests for materials should be addressed to S.J.M.

## METHODS

### Mouse studies and xenograft assays

Mouse experiments complied with all relevant ethical regulations and were performed according to protocols approved by the Institutional Animal Care and Use Committee at UT Southwestern Medical Center (protocol 2016-101360). For all experiments, mice were kept on normal chow and fed ad-libitum. Melanoma cell suspensions were injected subcutaneously in staining medium (L15 medium plus bovine serum albumin (1 mg/ml), 1% penicillin/streptomycin, and 10 mM HEPES (pH 7.4) with 25% high-protein Matrigel (product 354248; BD Biosciences)).

Melanoma specimens were obtained with informed consent from all patients according to protocols approved by the Institutional Review Board (IRB) of the University of Michigan (IRBMED approvals HUM00050754 and HUM00050085; see ref^69^) and UT Southwestern Medical Center (IRB approval 102010-051). Patient-derived melanomas were transplanted into 4 to 8-week-old male and female NOD.CB17-*Prkdc^scid^ Il2rg^tm1Wjl^*/SzJ (NSG) mice. Mouse melanomas were transplanted into 4 to 8-week-old NSG mice or 6 to 8-week-old male and female C57BL/Ka mice. Subcutaneous injections were performed in the right flank of mice in a final volume of 50 μl with 100 mouse or human cells per injection in NSG mice and 2 x 10^5^ mouse cells per injection in C57BL/Ka mice, unless otherwise specified. Subcutaneous tumor diameters were measured weekly with calipers until any tumor in the cohort reached 2.5 cm in its largest diameter. At that point, all mice in the cohort were killed, per approved protocol, for analysis of subcutaneous tumor diameter and metastatic disease burden. Metastatic disease burden was evaluated by bioluminescence imaging of visceral organs (see details below).

Unless otherwise specified, intravenous injections were performed by injecting 100 mouse melanoma cells or 1000 human melanoma cells into the tail veins of NSG mice or 10^5^ mouse melanoma cells into the tail veins of C57BL mice in 100 µl of PBS.

To deplete macrophages, mice were treated twice weekly with intraperitoneal injections of 200 μg endotoxin-free anti-CSF1R blocking antibody (BioXcell #BE0213) or an equal amount of endotoxin-free rat IgG2a isotype control antibody (BioXcell #BE0089). For experiments in which mice were depleted of macrophages before being injected with melanoma cells, mice were treated with anti-CSF1R or isotype control antibodies for two weeks prior to injection.

To treat mice with STING inhibitor (C-176, Med Chem Express HY-112906), mice received daily intraperitoneal injections with 10.8 mg/kg C-176 in corn oil (Sigma C8267), beginning when tumors became palpable. To treat mice with reverse transcriptase inhibitors, mice were gavaged daily with 60 mg/kg emtricitabine (Fisher Scientific AC46207005) and 100 mg/kg tenofovir (Fisher Scientific AC461250250) dissolved in 5% DMSO/30% PEG-300/1% Tween80. These suspensions were sonicated for ∼20 minutes, until fully dissolved, before use.

## Cell culture

YUMM 1.7 (*Braf^V600E/+^; PTEN^-/-^; Cdkn2^-/-^*), YUMM3.3 (*Braf^V600E/+^; Cdkn2^-/-^*), and YUMM5.2 (Braf^V600E/+^ p53^-/-^) mouse melanomas^30^ were obtained from, and authenticated by, the American Type Culture Collection (ATCC). Cell lines were confirmed to be mycoplasma free using the MycoAlert detection kit (product LT07-318, Lonza). YUMM1.7, YUMM 3.3, and YUMM5.2 were transfected with dsRed2 and luciferase (dsRed2-P2A-Luc) so that the cells could be identified and quantitated by flow cytometry and bioluminescence imaging. The cells were cultured in DMEM high-glucose (product 11965118, Thermo Fisher Scientific) supplemented with 10% Fetal Bovine Serum (FBS) and 1% penicillin/streptomycin. For injection into mice, cells were trypsinized using Trypsin-EDTA (0.25%; product 25300054, Thermo Fisher Scientific) for 2 to 3 minutes, then dissociated with gentle trituration to make single cell suspensions.

To perform spheroid formation assays, single cells from culture or from subcutaneous tumors were sorted, 1 cell per well, into 96-well Corning^TM^ Costar^TM^ Ultra-low attachment plates (Fisher Scientific 07-200-603) containing DMEM high-glucose (product 11965118, Thermo Fisher Scientific) supplemented with 10% FBS and 1% penicillin/streptomycin. Two to three weeks later, the percentage of wells that contained spheroids was assessed by microscopy. To analyze spheroids by flow cytometry, spheroids were gently mechanically dissociated then filtered through a 40-µm mesh to obtain single-cell suspensions.

## Phagocytosis assays

Phagocytosis assays were performed in culture using bone marrow-derived macrophages as described in^70^. Briefly, bone marrow from C57BL mice was plated in culture with 50 ng/ml M-CSF. Fresh M-CSF was added every two days. On day 7 of culture, M-CSF was removed and the macrophages were stimulated for 24 hours with 20 ng/ml IFN-y (Biolegend 575306) and 40 ng/ml lipopolysaccharide (Sigma L2880). Approximately 24 hours later, flow cytometrically-isolated melanoma cells were stained with Carboxyfluorescein succinimidyl ester (CFSE, BioTracker Green Cell Proliferation Kit from EMD Millipore SCT110) according to the manufacturer’s instructions. An equal number of macrophages and melanoma cells were co-cultured in DMEM high-glucose (product 11965118, Thermo Fisher Scientific) supplemented with 10% FBS and 1% pen/strep for 16-20 hours, then, the cells were analyzed by flow cytometry to determine the percentage of F4/80^+^ macrophages (PE-Cy7, Tonobo 60-4801-U025) that were CFSE^+^. 4’,6-Diamidino-2-phenylindole (DAPI; Invitrogen D1306) was added to exclude dead cells during flow cytometric analyses.

For *in vivo* phagocytosis assays, melanoma cells were isolated from subcutaneous tumors by flow cytometry, then stained with CFSE and injected intravenously into C57BL mice (2.5 x 10^5^ cells per injection). The next day, mice were killed and the lungs were chopped up and enzymatically digested with 0.26 U/ml Liberase^TM^ TL (Roche 05401020001) with 100 U/ml DNase for one hour at 37^°^C with agitation. Cells were then filtered through a 40-µm cell strainer to obtain single cell suspensions. Red blood cells were lysed by resuspending these cell suspensions in ammonium chloride potassium buffer (0.15 M ammonium chloride (Sigma A9434) plus 0.1 M potassium bicarbonate (Sigma 237205) and 0.1 mM ethylenediamine tetraacetic acid (EDTA, Fisher scientific BP2482) and incubating for 2-5 minutes at room temperature. After washing with HBSS, the cells were stained with antibodies to identify immune cells. In all cases, cells were gated on forward scatter height (FSC-H) versus forward scatter area (FSC-A) to exclude cell clumps, and DAPI was used to exclude dead cells. Mouse endothelial cells and non-lysed red blood cells were excluded by gating out cells that stained positively for CD31 or Ter119. All immune cells were identified as CD45^+^ as well as having the following markers: dendritic cells (CD24^+^CD11c^+^), macrophages (CD11b^+^F4/80^+^CD24^-^), NK cells (CD11b^+^F4/80^-^NK1.1^+^), monocytes (CD11b^+^Ly6C^+^Ly6G^-^CD115^+^CD11c^-^ from lung), T cells (CD3^+^CD11b^-^B220), B cells (B220^+^CD11b^-^CD3^-^), Tregs (CD3^+^CD4^+^FOXP3^+^CD11b^-^B220^-^) and neutrophils (CD11b^+^Ly6C^+^Ly6G^+^). The flow cytometry gates used to identify each cell population are shown in Extended Data Figure 4a.

## Bioluminescence imaging

Metastatic disease burden was assessed by bioluminescence imaging (all melanomas stably expressed luciferase). Five minutes before performing bioluminescence imaging, mice were injected intraperitoneally with 100 μl of PBS containing D-luciferin monopotassium salt (40 mg/ml; Biosynth, L8220), then mice were anaesthetized with isoflurane 2 minutes prior to imaging. The mice were imaged using an IVIS Imaging System 200 Series (Caliper Life Sciences) with Living Image software (Perkin Elmer). After whole-body imaging, the visceral organs were surgically removed and imaged. The exposure time was set to ‘auto’, and ranged from 10 to 60 s, depending on the maximum signal intensity, to avoid saturation of the luminescence signal. To measure background luminescence, a negative control mouse not transplanted with melanoma cells was imaged. The bioluminescence signal (total photon flux) was quantified with ‘region of interest’ measurement tools in Living Image software. Metastatic disease burden was calculated as observed total photon flux across all organs minus background total photon flux in negative control mice not injected with melanoma cells. Data were then normalized to the control mice as indicated in figure legends.

## Isolation and analysis of melanoma cells by flow cytometry

To isolate melanoma cells from circulation, mice were transplanted subcutaneously with human melanoma cells and the tumors were allowed to grow until they reached a maximum of 2.5 cm. Approximately 500 µL of blood was collected by cardiac puncture and added to 150 µL of citrate-phosphate-dextrose solution (Sigma C7165) to prevent clotting. Next, the blood samples were sedimented using Ficoll according to the manufacturer’s instructions (product 17144002, Ficoll Paque Plus, GE Healthcare) to eliminate red blood cells. The cells that remained in suspension were washed with Hank’s Balanced Buffered Solution (product 14025076, HBSS, Gibco) and then stained with antibodies for flow cytometric analysis.

To isolate melanoma cells from subcutaneous or metastatic tumors, mice were transplanted subcutaneously with human or mouse melanoma cells and the cells were allowed to spontaneously metastasize until the subcutaneous tumor reached a maximum of 2.5 cm in diameter. The tumors were then removed, chopped up and enzymatically dissociated in 200 U/ml collagenase IV (Worthington) for 20 minutes at 37^°^C. DNase (50-100 U/ml) was added to reduce cell clumps during digestion. Cells were then filtered through a 40-µm cell strainer to obtain single-cell suspensions.

All antibody staining was performed for 20 minutes on ice, followed by washing with Hank’s Buffered Salt Solution (HBBS) and centrifugation at 200xg for 5 min. Cells were stained with directly conjugated antibodies against mouse CD45 (PE-Cy7, Tonbo 60-0453-U100, violet Fluor 450, Tonbo 75-0451-U100, APC, Tonbo 20-0451-U100), mouse CD31 (PE-Cy7, Biolegend 102418, violet Fluor 450, Invitrogen 48-0311-82 or APC, Biolegend 102410), mouse Ter119 (PE-Cy7, Biolegend 116223, violet Fluor 450, Tonbo 75-5921-U100 or APC, Biolegend 116212) and human HLA-A, B, C (FITC, BD Pharminigen 555552). Human melanoma cells were isolated as cells that were positive for HLA and DsRed as well as negative for mouse endothelial and hematopoietic markers. The flow cytometry gates used to identify human melanoma cells are shown in Extended Data Figure 1b. Mouse melanoma cells were isolated as cells that were positive for DsRed and Melanoma Cell Adhesion Molecule (MCAM) (APC, BioLegend 134712 or PE-Cy7, Biolegend 134713) and negative for mouse CD45, Ter119, and CD31. The flow cytometry gates used to identify mouse melanoma cells are shown in Extended Data Figure 1c. Cells were washed with HBSS and resuspended in 4’,6-diamidino-2-phenylindole (DAPI; 1 μg/ml; Sigma) to eliminate dead cells from sorts and analyses. Cells were analyzed using either a FACS Fusion, FACS Aria II SORP or FACS Lyric (BD Biosciences). Prior to re-injection of sorted cells, purity was confirmed by post-sort analysis of sorted cell populations. Cells were counted using a hemocytometer (trypan blue exclusion) and resuspended in HBSS for intravenous injections or staining medium for subcutaneous injections.

To measure the levels of CD47, PD-L1, B2M and H2-Kb on mouse melanoma cells, single cell suspensions from subcutaneous or metastatic tumors (generated as detailed above) were stained with the following antibodies: CD47 (BioLegend 127514), PD-L1 (Tonbo 20-1243-U100), H2-Kb (BioLegend 116518), and B2M (BioLegend 154506). Background was determined by staining cells with mouse IgG2a (BioLegend 400220) or rat IgG2a (BioLegend 400512) or isotype control antibodies. All antibody cocktails also contained mouse lineage markers (CD45, Ter119, CD31) to exclude mouse hematopoietic and endothelial cells.

For antibody blocking experiments, sorted cells were counted and treated with endotoxin-free antibodies against PD-L1 (SelleckChem number A2115), CD47 (BioXcell number BE0270), trinitrophenol (rat IgG2a isotype control, BioXcell number BE0089), or keyhole limpet hemocyanin (rat IgG2b isotype control, BioXcell number BE0090) for 30 minutes on ice.

Unbound antibodies were then washed off and cells were resuspended in HBSS before intravenous injection. To isolate immune cells for qRT-PCR analysis, tumors were digested as detailed above and 10,000-50,000 cells were sorted into RLT buffer (Qiagen RNeasy Micro Kit) for RNA purification according to the manufacturer’s instructions. Antibodies against the following markers were used to identify immune cells: CD45 (PE-Cy7 or PE, Biolegend 103106), CD11b (APC-780, eBiosciences 47-0112-82), Ter119 (APC, Biolegend 116212 or FITC, Tonbo 35-5921-U100), F4/80 (BioLegend 123116), CD11c (FITC, BioLegend 117306), CD24 (PE, BioLegend 114308), CD115 (BV510, Biolegend 750893), CD88 (APC, Biolegend 135808), CD3 (APC-Cy7, Biolegend 100222), CD8a (Brilliant Violet 510, BioLegend 100738), CD4 (BV421, BioLegend 100559), NK1.1 (AF700, BioLegend 156512), FOXP3 (AF700, BioLegend 126422), Ly6C (BV421, BD 562727), Ly6G (PE, Biolegend 164504), TCRy/o (APC, BioLegend 111206). Fixed and permeabilized cells were stained with anti-FOXP3 antibody to identify regulatory T cells (BD product number BDB552598). The flow cytometry gates and marker combinations used to identify each immune cell population are shown in Extended Data Figure 4a.

## Immunofluorescence analysis of tumor sections

Tumors were harvested from C57BL mice when they reached 1.5-2.0 cm in diameter, then fixed in 4% paraformaldehyde (Fisher Scientific product AAJ19943K2) for 24-48 hours at 4°C. The tumors were then washed, cryopreserved in 30% sucrose for ∼48 hours, embedded in Optimal Cutting Temperature (OCT) compound, and frozen. 8-12 µm sections were cut, then stained with antibodies against F4/80 (Thermo Scientific 14-4801-82; 1:200) to identify macrophages, DsRed (LifeSpan Biosciences LS-C340696, 1:500) to identify melanoma cells, and GFP (Aves Labs GFP-1020; 1:500 or Thermo Scientific A-11122; 1:500) to identify GFP-ISG15^+^ cells. Melanoma specimens from patients were fixed in 10% formalin overnight, washed, then cryopreserved and sectioned as described above, and stained with antibodies against Melanoma Cell Adhesion Molecule (MCAM, Novus Biologicals NBP2-44512; 1:200) to identify melanoma cells, ISG15 (Abcam ab227451; 1:100), CD68 to identify macrophages/monocytes (Abcam ab201340; 1:100) and/or IFNα (Thermo Scientific PA5-119649; 1:200). Images were collected using a Nikon W1 SoRa Spinning disk confocal microscope and processed in Fiji (ImageJ 1.53q) or Imaris (Oxford Instruments) software. To measure the distance between GFP-ISG15^-^ or GFP-ISG15^+^ cells and macrophages in tumors, Imaris was used to generate surfaces for each cell type and distances were determined using Imaris.

## qRT-PCR analyses

To compare *ISG15, IFI27* and *IFITM3* transcript levels in human melanoma cells, the melanoma cells were isolated by flow cytometry from subcutaneous tumors, the blood, or metastatic tumors of the same NSG mice bearing 2.0-2.5 cm subcutaneous tumors using the methods detailed above. To assess *IFN*5*1* transcript levels in immune cells, immune cells were sorted from subcutaneous tumors grown in C57BL mice. In all cases, RNA was extracted from the cells using an RNeasy Mini Kit (product 74106, Qiagen) according to the manufacturer’s instructions. RNA was reverse transcribed using SuperScript III (product 18080044, Invitrogen). qRT-PCR was performed using a Roche LightCycler 480. The primers used for qRT-PCR analysis of RNA were h*ISG15*-Fwd GACATTCGGCTGTTTACC and h*ISG15*-Rev GCGGTTCTGTGGAGGTTA; h*IFI27*-Fwd CGTCCTCCATAGCAGCCAAGAT and h*IFI27-*Rev ACCCAATGGAGCCCAGGATGAA; *hIFITM3-*Fwd TGGCCAGCCCCCCAACTAT and h*IFITM3*-Rev CATAGGCCTGGAAGATCAG; m*IFNb*-Fwd TCCGAGCAGAGATCTTCAGGAA and m*IFNb-*Rev TGCAACCACCACTCATTCTGAG. mIFNa-Fwd: GGACTTTGGATTCCCGCAGGAGAAG; Rev: GCTGCATCAGACAGCCTTGCAGGTC.

## Bulk RNA sequencing

To isolate melanoma cells from patient-derived xenografts for RNA sequencing, tumors were allowed to grow to 1.5-2.0 cm in NSG mice. Mice were then killed, and cells were isolated from subcutaneous tumors, the blood or metastatic nodules by flow cytometry as described above. To isolate GFP-ISG15^-^ or GFP-ISG15^+^ cells for RNA sequencing, mouse melanoma tumors were allowed to grow to 1.5-2.0 cm in C57BL mice. For YUMM1.7, three spontaneous lymph node metastatic tumors were also included. Tumors were harvested and cells were isolated by flow cytometry as detailed above. Cells were then lysed and RNA was extracted using RNeasy Mini Kits (product 74106, Qiagen). RNA quality was confirmed using RNA ScreenTapes (Agilent Technologies 5067-5582). Libraries were generated using SMARTer Stranded Total RNA-Seq kit – v2 Pico Input Mammalian (Takara). Library fragment size was measured using D1000 Screen Tape (Agilent) and libraries were quantified using the Qubit dsDNA high-sensitivity assay kit (Life Technologies).

Samples were sequenced on an Illumina HiSeq 2000 or NextSeq 2000 platform with 100 bp single or paired-end read settings. Data were analyzed with the bulk RNA-seq workflow developed by the Morrison Lab (https://git.biohpc.swmed.edu/CRI/morrison-lab/RNASeq). Raw read quality was assessed using FastQC (0.12.1). Raw reads were trimmed using Trim Galore (0.6.4) and were aligned to the Ensembl GRCh38 human or GRCm38 mouse reference genomes using STAR (2.7.9a). Mapped reads were quantified using HTSeq-Count (2.0.9) with the “--nonunique all option” or TEcount from TEtranscripts (2.2.1) with the “–mode multi option”. Exon-mapped reads were normalized, and gene expression levels were measured as fragments per thousand exonic bases per million mapped reads (FPKMs) using DESeq2 (1.50.2) with R (4.5.2). Differential expression tests were performed using DESeq2 as well.

Gene Ontology Biological Process (GOBP) over-representation analysis was performed with clusterProfiler (4.16.0) with R (4.5.2). Differentially expressed (DE) genes were defined as those with False Discovery Rate (FDR) <0.05 and fold change >1.5. Prior to over-representation analysis, a predefined set of cell cycle genes (Seurat S and G2/M markers derived from ref^71^) were removed from the differentially expressed gene list to limit enrichment of gene sets based on cell cycle genes. Gene Set Enrichment Analysis (GSEA) was performed on the log2-transformed fold change values using GSEAPreranked from GSEA (4.1.0) with a combined database including the MSigDB (v2023.1) gene sets and the Transposable Element (TE) gene sets from TEtranscripts. The per-sample Gene Set Variation Analysis (GSVA) scores were calculated from log2-transformed FPKM values using the Gaussian kernel estimation method of GSVA (2.5.15) with R (4.5.2). RNA-seq data were stored, processed, and analyzed using the high-performance computing core (BioHPC) at UT Southwestern Medical Center.

## Gene targeting in mouse melanomas

GFP-ISG15 and GFP-IFIT3 knock-in reporters were generated using the CRISPR TrueTag^TM^ Donor DNA Kit, GFP (product A53806, Thermo Scientific). Cells were transfected according to the manufacturer’s protocol using Lipofectamine^TM^ CRISPRMAX^TM^ Cas9 Transfection reagent and TrueCut^TM^ Cas9 Protein v2 (products CMAX00003 and A36498, respectively, Thermo Scientific). gRNAs were designed using Thermo Scientific’s genome editor tool. For all knock-ins, only the N-terminal gRNAs were used: mouse *ISG15* gRNA (5’—CAGCAATGGTGAGTCCAATT—3’), mouse *IFIT3* gRNA (5’—AGATTGCACTGACCTCATGA—3’) were used with donor DNA generated according to the manufacturer’s protocol. Primers used to generate donor DNA were: mouse *ISG15* forward (5’—TTCCAGGGGACCTAGAGCTAGAGCCTGCAGCAATGGGAGGTAAGCCCTTGCATTCG—3’) and reverse (5’—AACTAAAATGAAACCCATCTGCCTCTGTCCCAAATTGGACTCACACCGCTTCCACTACCTGA ACC—3’), mouse *IFIT3* forward (5’—GCTTTTCCCAGCAGCACAGAAACAGATCACCATCATGGGAGGTAAGCCCTTGCATTCG—3’) and reverse (5’—TCATTTAATTGGAGAAGCAGAGATTGCACTGACCTACCGCTTCCACTACCTGAACC—3’).

Since expression of puromycin resistance in edited cells was regulated by endogenous *ISG15* or *IFIT3* promoters, the transfected cells were first stimulated overnight with 10 ng/mL IFN51 (Biolegend 581304) approximately 48 hours after transfection. After 48 hours of IFN51 stimulation to induce *ISG15* or *IFIT3* expression, cells were treated with puromycin (product number 4089, Tocris) to select for edited cells. After selection, cells were removed from IFN51/puromycin for 7 days and then restimulated with IFN51 overnight to induce reporter expression. Cells that expressed GFP in the presence of IFN51 were sorted, one cell per well, into 96-well plates. Clones were then screened by PCR and Sanger sequencing to ensure 5’ and 3’ ends were inserted on target and in frame. Finally, clones were again confirmed to selectively express GFP upon treatment with IFN51.

To make *IFNAR1*, *CD47* and *B2M* mutant mouse melanomas, a TrueTag^TM^ Knockout Enrichment Donor DNA Kit (Thermo Scientific product number A53815) was used to insert stop codons immediately after the start codon in the first exon of the targeted gene to prevent translation. In each gene, we found no alternative, in-frame start codons in subsequent exons. Donor DNA was synthesized according to the manufacturer’s instructions. Donor DNA contained a puromycin resistance gene cassette used to select for cells with the donor DNA, followed by red fluorescence protein (RFP) and three consecutive stop codons to prevent translation of the targeted protein. Although this RFP could not be distinguished from the DsRed tags in the same cell lines, this did not matter as the DsRed/RFP signal was only used to distinguish melanoma cells from non-melanoma cells.

Cells were transfected with the gRNA, Cas9 protein and donor DNA according to the manufacturer’s instructions. Control cells were transfected with Cas9 protein and a scrambled sgRNA. After transfection, cells were selected with puromycin for one week (control cells were not treated with puromycin) and then stained with antibodies against the targeted protein to confirm deficiency. Clones were screened by first confirming the absence of the target protein by flow cytometry (IFNAR1, APC, Biolegend 127314, the antibodies used for CD47, B2M and H2-Kb are listed above) followed by Sanger sequencing to confirm on-target insertion of the donor DNA. gRNA sequences were as following: *IFNAR1* #1 CAAGACGATGCTCGCTGTCG and #2 AAGACGATGCTCGCTGTCGT. *CD47* #1 TTGCATCGTCCGTAATGTGG and #2 CCCTTGCATCGTCCGTAATG. *B2M* #1 CATGGCTCGCTCGGTGACCC and #2 AGTCGTCAGCATGGCTCGCT. *IFNAR1* homology arm primers, #1 Fwd AGCCGCCGCCCGGCCTCCCAAGACGATGCTCGCTGTCGGAAGTGGCTCAGGTTCTGGA and rev GCCCCGGCCACCAGCACCAGGGCCGCCGCGCCCACCTTGGCCGATCGCATACAGAG. #2 Fwd GCCGCCGCCCGGCCTCCCAAGACGATGCTCGCTGTCGGAAGTGGCTCAGGTTCTGGA and rev GCCCCGGCCACCAGCACCAGGGCCGCCGCGCCCACCTTGGCCGATCGCATACAGAG.

*CD47* homology arm primers #1 Fwd

TGAAACTGTGGTCATCCCTTGCATCGTCCGTAATGTGGGAAGTGGCTCAGGTTCTGGA and Rev CACTTCACAAACATTTCTTCGGTGCTTTGCGCCTCCTTGGCCGATCGCATACAGAG. #2 Fwd CAATGAAACTGTGGTCATCCCTTGCATCGTCCGTAATGGAAGTGGCTCAGGTTCTGGA and Rev TTCACAAACATTTCTTCGGTGCTTTGCGCCTCCACCTTGGCCGATCGCATACAGAG. *B2M*

homology arm primers #1 Fwd GGTCGCTTCAGTCGTCAGCATGGCTCGCTCGGTGACCGGAAGTGGCTCAGGTTCTGGA

and Rev AGGCCGGTCAGTGAGACAAGCACCAGAAAGACCAGCTTGGCCGATCGCATACAGAG. #2

Fwd TTCAGTCGCGGTCGCTTCAGTCGTCAGCATGGCTCGCGGAAGTGGCTCAGGTTCTGGA

and Rev AGTGAGACAAGCACCAGAAAGACCAGGGTCACCGACTTGGCCGATCGCATACAGAG.

## Statistical methods

In each type of experiment, multiple melanomas were tested in multiple independent experiments, usually performed on different days. Mice were allocated to experiments randomly and samples processed in an arbitrary order, but formal randomization techniques were not used. No formal blinding was applied when performing the experiments or analyzing the data.

Prior to analyzing the statistical significance of differences among treatments, we tested whether the data were normally distributed and whether variance was similar among treatments. To test for normal distribution, we performed the Shapiro–Wilk test when 3≤n<20 or the D’Agostino Omnibus test when n≥20. To test if variability significantly differed among treatments we performed *F*-tests (for experiments with two treatments) or Levene’s median tests (for more than two treatments). When the data significantly deviated from normality or variability significantly differed among treatments, we log2-transformed the data and tested again for normality and variability. Fold change data were always log2-transformed. If the transformed data no longer significantly deviated from normality and equal variability, we performed parametric tests on the transformed data. If log2-transformation was not possible or the transformed data still significantly deviated from normality or equal variability, we performed non-parametric tests on the non-transformed data.

All of the statistical tests we used were two-sided, where applicable. To assess the statistical significance of a difference between two treatments, we used Student’s *t*-tests (when a parametric test was appropriate), Welch’s *t*-tests (when a parametric test was appropriate after correcting data variances), or Mann-Whitney tests (when a non-parametric test was appropriate). Multiple *t*-tests (parametric or non-parametric) were followed by Holm-Sidak’s multiple comparisons adjustments. To assess the statistical significance of differences between more than two treatments, we used ordinary or matched samples one-way or two-way ANOVAs (when a parametric test was appropriate) or mixed effects analyses (when a parametric test was appropriate and there were missing data points) followed by Tukey’s, Dunnet’s or Sidak’s multiple comparisons adjustments or Kruskal-Wallis tests (when a non-parametric test was appropriate) followed by Dunn’s multiple comparisons adjustment. Multiple comparisons were adjusted per melanoma cell line as a family where applicable, or among multiple cell lines when there were two treatments per cell line.

To assess the statistical significance of differences between time-course data, we used nparLD^72^, a statistical tool for the analysis of non-parametric longitudinal data, followed by the Benjamini-Hochberg False Discovery Rate (FDR) method for multiple comparisons adjustment. All statistical analyses were performed with Graphpad Prism 10.6.1 or R 4.5.2 with the stats, fBasics, car, and nparLD packages. All data represent mean ± standard deviation.

Samples sizes were not pre-determined based on statistical power calculations but were based on our experience with these assays. For assays in which variability is commonly high, we typically used n>10. For assays in which variability is commonly low, we typically used n<10.

No data were excluded; however, mice sometimes died during experiments, presumably due to the growth of metastatic tumors. In those instances, data that had already been collected on the mice in interim analyses were included (such as subcutaneous tumor growth measurements over time) even if it was not possible to perform the end-point analysis of metastatic disease burden due to the premature death of the mice.

## Data availability

Source Data files contain the numeric data for all figures and extended data figures. Other data are available from the corresponding author upon reasonable request. Bulk RNA sequencing data are available from the National Center for Biotechnology Information (NCBI) BioProject ID PRJNA1433150 (comparing the gene expression profiles of GFP-ISG15^-^ versus GFP-ISG15^+^ mouse melanoma cells) and PRJNA1440044 (comparing the gene expression profiles of melanoma cells from primary subcutaneous tumors, the blood, and metastatic tumors in patient-derived xenografts).

---

**Extended Data Figure 1.**
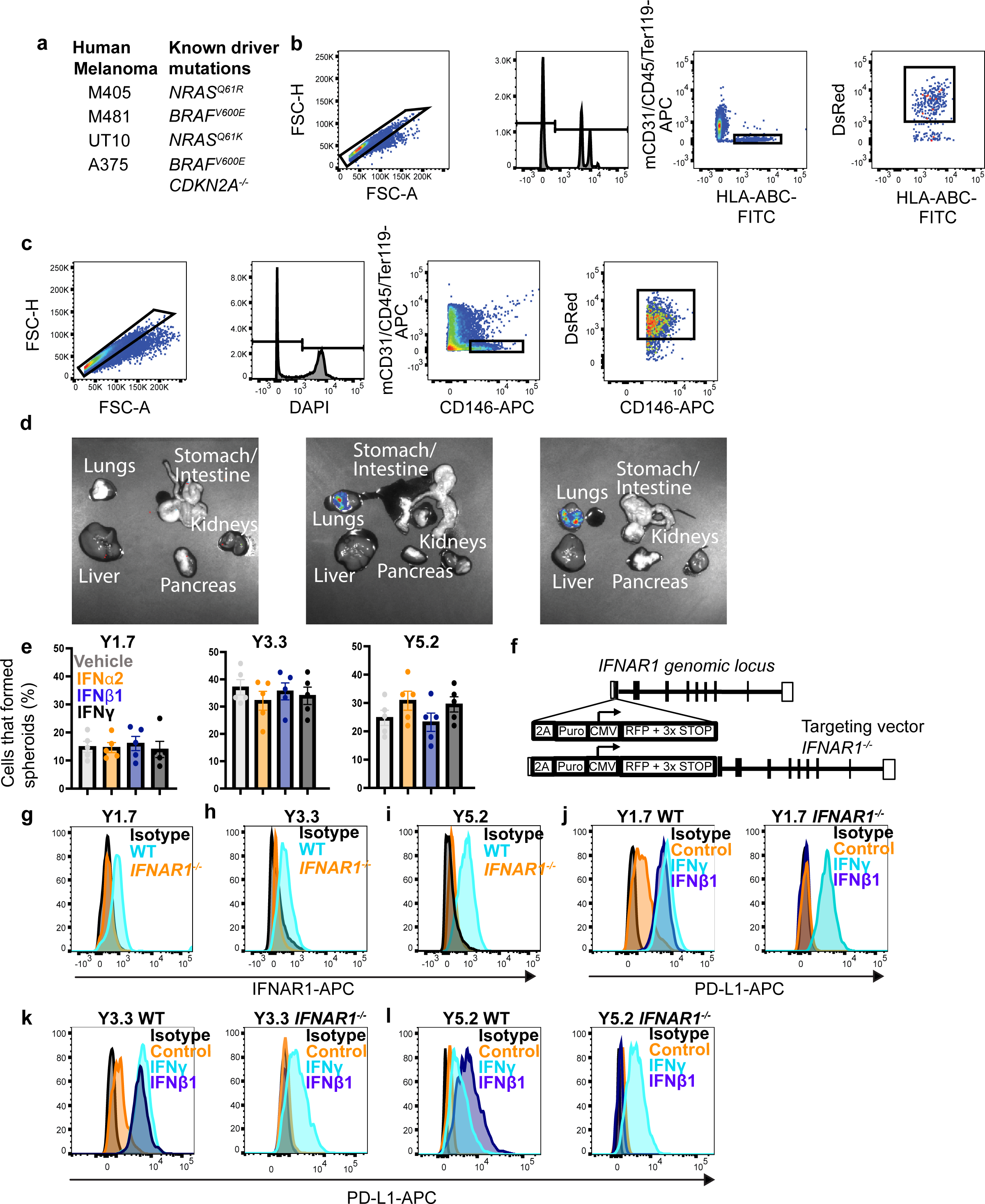
Representative flow cytometry gating strategies for the isolation of melanoma cells, representative bioluminescence imaging of metastatic disease, *IFNAR1* targeting strategy, and validation of IFNAR deficiency in targeted clones. Related to figures 1 and 2. a, Human melanomas that were used in Figure 1 along with their known driver mutations. b-c, Flow cytometry plots showing the gating strategies used to identify human (b) or mouse (c) melanoma cells. In all cases, cells were gated on forward scatter height (FSC-H) versus forward scatter area (FSC-A) to exclude cell clumps, DAPI staining was used to exclude dead cells, and mouse hematopoietic and endothelial cells were excluded by gating out cells that stained positively for anti-mouse CD45, CD31, and/or Ter119. Human melanoma cells were selected by including cells that stained positively for HLA-ABC and DsRed, which was stably expressed in all melanomas along with luciferase. Mouse melanoma cells were selecting by including cells that stained positively for Melanoma Cell Adhesion Molecule (MCAM; CD146) and DsRed. d, Representative bioluminescence imaging of visceral organs dissected from a negative control mouse (left) or a mouse injected intravenously with human (center) or mouse (right) melanoma cells. The presence of melanoma cells in visceral organs is indicated by blue/green/red signal and measured as total photon flux (photons/s). For each experiment, a control mouse not injected with melanoma cells was analyzed to determine background signal, which was subtracted from each of the mice injected with melanoma cells. To combine data from multiple experiments, data were normalized to mice injected with control cells. e, Percentage of melanoma cells cultured overnight in the indicated interferons that formed spheroids upon subcloning into secondary cultures (each dot represents a different 96-well plate; two experiments with a total of five plates per melanoma). f, The targeting strategy used to generate *IFNAR1* mutant melanomas. (g-i) Representative flow cytometry histograms showing loss of anti-IFNAR1 antibody staining in I*FNAR1*-mutant mouse melanoma cells (histograms are representative of three independently-targeted *IFNAR1* mutant clones per melanoma). (j-l) Loss of IFNAR1 activity: control cells exhibited increased PD-L1 levels in response to culture in interferon y and interferon 51 but *IFNAR1* mutant cells exhibited increased PD-L1 lin response to interferon y but not interferon 51 (histograms are representative of three independently-targeted *IFNAR1* mutant clones per melanoma). All data represent mean ± s.d. Statistical significance was assessed using a two-way ANOVA followed by Dunnett’s multiple comparisons adjustment (e). All statistical tests were two-sided. No statistically significant differences were observed in panel e.

**Extended Data Figure 2.**
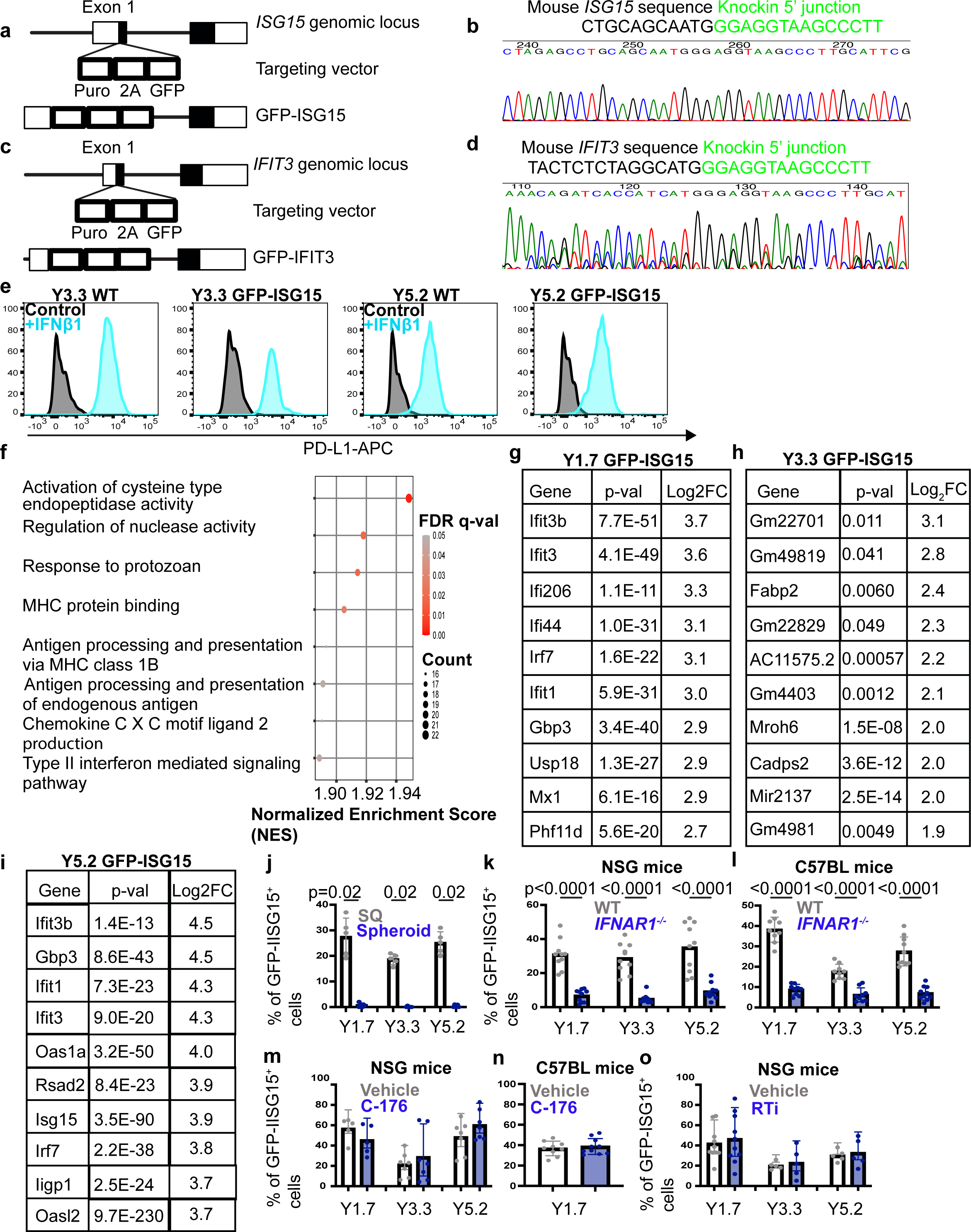
Generation of GFP-ISG15 and GFP-IFIT3 knock-in reporters. Related to. figure 2**. a-d**, Targeting strategies (**a, c**) for the generation of GFP-ISG15 and GFP-IFIT3 fusion protein reporters in which GFP was inserted into the first exon of both interferon-regulated genes. Correctly targeted alleles were confirmed by Sanger sequencing of the junction sites (**b, d**). Endogenous *ISG15/IFIT3* sequences are shown in black and inserted GFP sequences are shown in green (**b, d**). Each reporter-bearing melanoma clone was confirmed to be heterozygous for the indicated reporter. **e**, Representative flow cytometry plots showing induction of PD-L1 in control (WT) or GFP-ISG15 mouse melanoma cells treated with interferon 51 in culture. **f-i**, Bulk sequencing of RNA from GFP-ISG15^-^ and GFP-ISG15^+^ melanoma cells that were flow cytometrically isolated from subcutaneous tumors grown in C57BL mice. The most significantly enriched gene sets in YUMM1.7 GFP-ISG15^+^ as compared to GFP-ISG15^-^cells are shown in (**f**) (GSEA NES > 1.5, FDR <0.05). In **g-i**, the 10 most significantly enriched genes (FDR < 0.05) in GFP-ISG15^+^ as compared to GFP-ISG15^-^ cells are shown for each mouse melanoma. **j**, Unfractionated cells from GFP-ISG15 bearing mouse melanomas were added to culture and then the percentage of GFP-ISG15^+^ cells in the spheroids that formed in culture was plotted beside the percentage of GFP-ISG15^+^ cells in the subcutaneous tumors from which they derived. Few GFP-ISG15^+^ cells were observed in culture, consistent with a lack of interferon production by melanoma cells in culture (two experiments with a total of five mice per melanoma). **k-l**, The percentage of cells that were GFP-ISG15^+^ in subcutaneous tumors that formed in NSG (**k**) or C57BL (**l**) mice by *IFNAR1* mutant or control melanoma cells bearing the GFP-ISG15 reporter (two experiments with a total of 9-10 mice per melanoma). **m-n**, The percentage of cells that were GFP-ISG15^+^ in subcutaneous tumors in NSG (**m**) or C57BL (**n**) mice treated with STING inhibitor (C-176) or vehicle control (one to two experiments with a total of 5-10 mice per melanoma). **o**, The percentage of cells that were GFP-ISG15^+^ in subcutaneous tumors in NSG mice treated with reverse transcriptase inhibitors or vehicle control (one to twoexperiments with a total of 5 to 10 mice per melanoma). Each dot represents a different mouse in panels **j-o** and all data represent mean ± s.d. Statistical significance was assessed using Mann-Whitney tests followed by Holm-Sidak multiple comparisons adjustments (**j**), two-way ANOVAs followed by Sidak’s multiple comparisons adjustments (**k, m**, and **o**), Welch’s t-tests followed by Holm-Sidak’s multiple comparisons adjustment (**l**), or a Student’s *t*-test (**n**). All statistical tests were two-sided. No statistically significant differences were observed in (**m-o**).

**Extended Data Figure 3.**
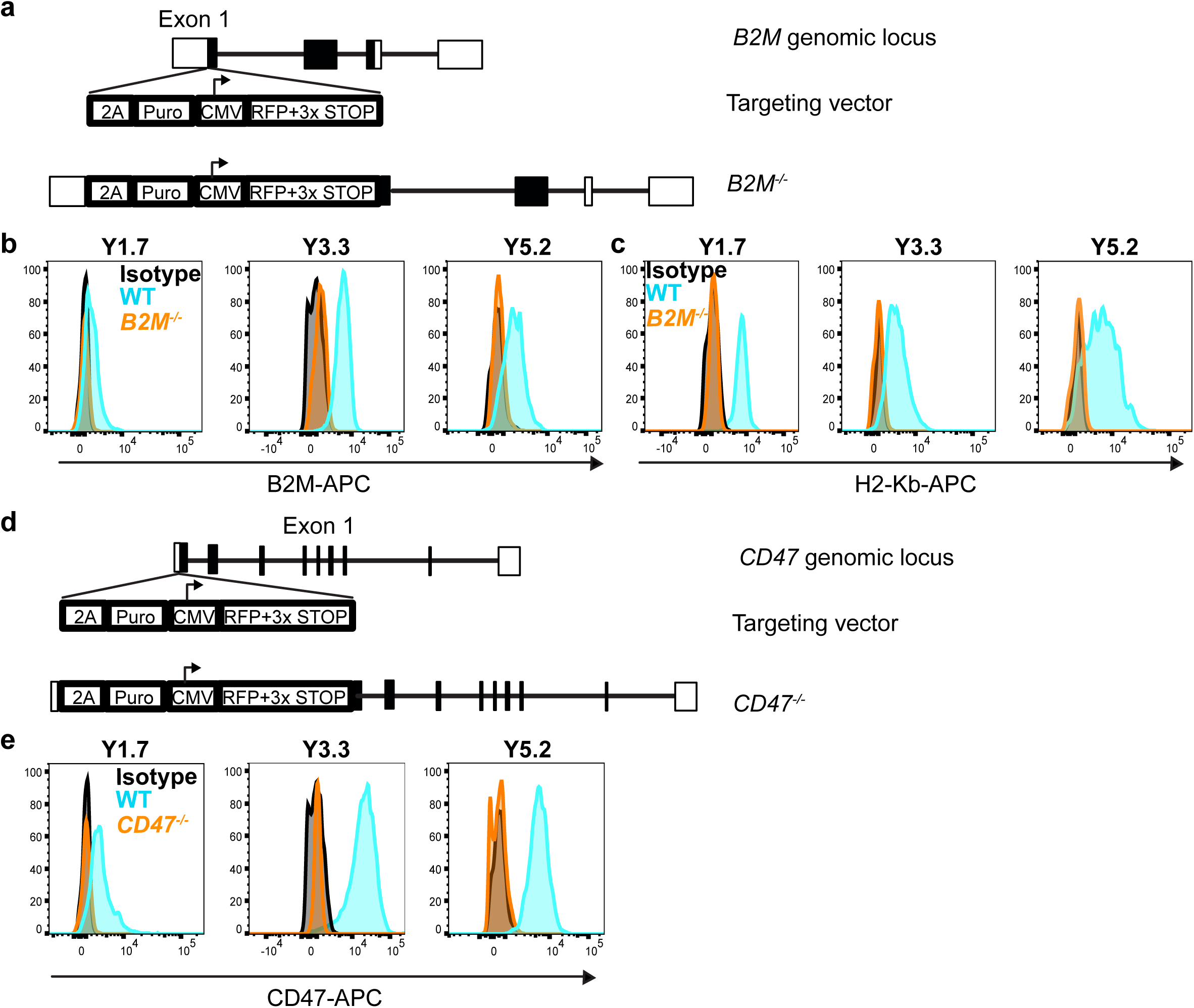
Targeting strategy and validation for *B2M* and *CD47* mutant melanomas. Related to. figure 3**. a**, Targeting strategy used to generate *B2M* mutant melanomas by knocking STOP codons into the first exon of *B2M* to prevent translation. **b**,**c** Representative flow cytometry histograms showing the lack of B2M or H2-Kb staining on the surface of *B2M* mutant melanomas. Since cells in culture have relatively low surface levels of B2M and H2-Kb, all clones were first cultured overnight with 10 ng/mL mIFN51 to increase cell surface levels of B2M and H2-Kb. One representative clone is shown for each of the *B2M* mutant YUMM lines (1.7, 3.3, and 5.2) that were generated. **d**, Targeting strategy used to generate *CD47* mutant melanomas by knocking STOP codons into the first exon of *CD47* to prevent translation. **e** Representative flow cytometry histograms showing the lack of CD47 staining on the surface of *CD47* mutant melanomas. All cells were first cultured overnight with 10 ng/mL mIFN51 to increase cell surface levels of CD47. One representative clone is shown for each of the *CD47* mutant YUMM lines (1.7, 3.3, and 5.2) that were generated. For each YUMM line, two control clones and three independently-targeted *B2M* mutant clones were used in further experiments.

**Extended Data Figure 4.**
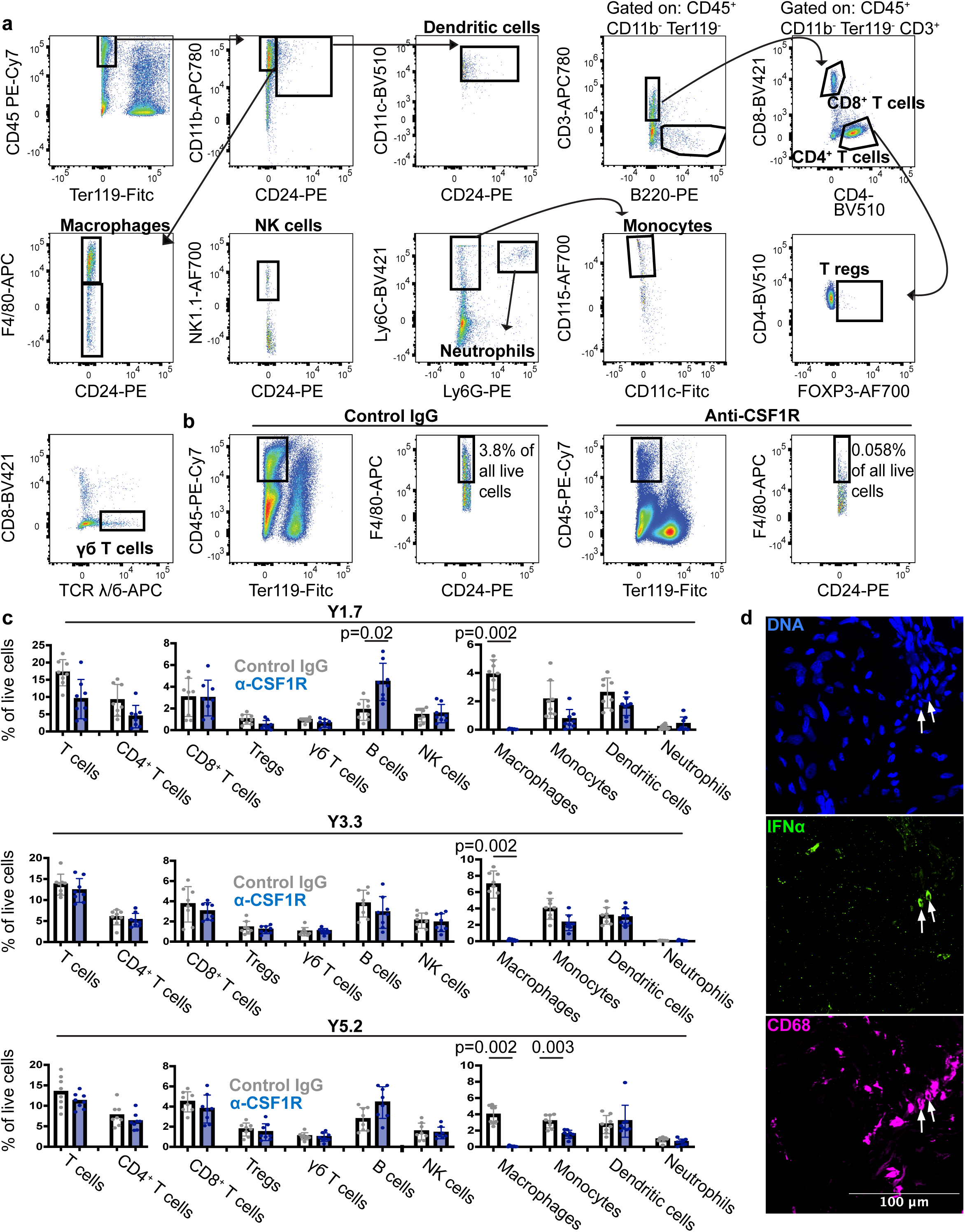
Markers and flow cytometry gating strategies used to identify immune cell populations and confirmation that treatment of mice with anti-CSF1R depleted macrophages/monocytes. Related to figures 4, 5, and 6. a**, Representative flow** cytometry plots showing the gating strategies used to identify mouse immune cells in primary tumors or lungs. In all cases, cells were gated on forward scatter height (FSC-H) versus forward scatter area (FSC-A) to exclude cell clumps and propidium iodide staining was used to exclude dead cells. All immune cells were first identified as being CD45^+^ and Ter119^-^, then further gated on the following markers to identify each cell population: dendritic cells (CD24^+^CD11c^+^), macrophages (CD11b^+^F4/80^+^CD24^-^), NK cells (CD11b^+^F4/80^-^NK1.1^+^), monocytes (CD11b^+^Ly6C^+^Ly6G^-^CD115^+^CD11c^-^ from lungs and CD11b^+^Ly6C^+^Ly6G^-^CD88^+^CD11c^-^ from tumors), T cells (CD3^+^CD11b^-^B220^-^), B cells (B220^+^CD11b^-^CD3^-^), Tregs (CD3^+^CD4^+^FOXP3^+^CD11b^-^B220^-^) and neutrophils (CD11b^+^Ly6C^+^Ly6G^+^). **b**, Consistent with prior studies^68^, treatment of mice with anti-CSF1R antibody depleted F4/80^+^ macrophages from tumors. **c**, Quantification of immune cell types in YUMM1.7 (top), YUMM3.3 (middle), or YUMM5.2 (bottom) subcutaneous melanomas in C57BL mice treated with anti-CSF1R or isotype control antibody. Each dot represents a different mouse (two experiments with a total of eight mice per melanoma). **d**, Melanoma specimens that were surgically resected from patients were stained with DAPI (blue) to label nuclei, anti-CD68 antibody (fuchsia) to label macrophages/monocytes and anti-IFNα antibody (data are representative of samples from two patient specimens). All data represent mean ± s.d.. Statistical significance was assessed using Student’s *t*-tests or Mann-Whitney tests followed by Holm-Sidak adjustments for multiple comparisons (**c**). All statistical tests were two-sided.

